# Ca^2+^-mediated protein citrullination regulates proliferation in the regenerating and malignant CNS

**DOI:** 10.64898/2026.01.25.701650

**Authors:** Samuel H. Crossman, Thamodi Walpola, Alon M. Douek, Jingyi Wang, Mitra Amiri Khabooshan, Connie Fung, Graham J. Lieschke, Iman Azimi, Jan Kaslin

**Affiliations:** Australian Regenerative Medicine Institute, Monash University; Monash University; Australian regenerative medicine institute, Monash University, Biomedicine Institute, University of Turku

**Keywords:** Stem Cells, Regeneration, Cancer, Spinal Cord Injury, dsRNA, TLR3, Inflammation, mechanoreception, *in vivo* imaging

## Abstract

Unlike most adult mammals, regenerative vertebrates can awaken dormant neural stem cells in response to injury. Understanding how this process is regulated could guide strategies to activate stem cells for tissue repair and limit aberrant proliferation in cancers of the CNS. Here, using zebrafish injury models and high-speed live imaging, we identify hydrodynamically-activated Ca²⁺ signalling as a key driver of neural stem cell activation. Local injury-associated changes in CSF flow activate mechanoreceptors at the site of spinal cord lesions, triggering pulsatile Ca²⁺ activity and progenitor proliferation. We identify Ca²⁺-regulated peptidylarginine deiminase enzymes (PADs) as key downstream effectors that citrullinate intracellular targets in a Ca²⁺-dependent manner to drive progenitor activation. Finally, we show that PAD inhibitors suppress the growth of aggressive medulloblastoma cells in preclinical laboratory models. Together, these findings uncover a novel mechanism of proliferation control in the vertebrate CNS and highlight the value of regenerative studies for identifying therapeutic targets.

**Graphical Summary:** 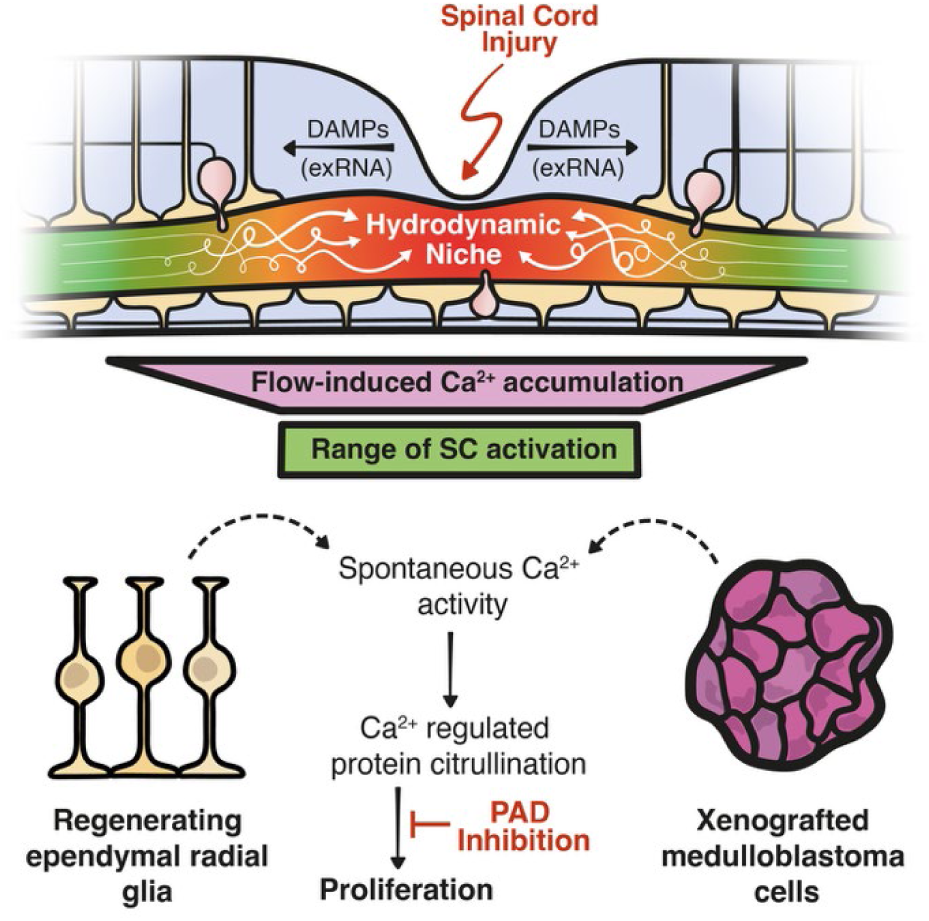

**Highlights:**

1. Damage signals trigger changes in cilia activity and CSF flow to create a transient hydrodynamic niche at the site of spinal cord injuries.
2. Specialised CSF-contacting neurons detect altered CSF circulation and initiate a signalling relay that culminates in elevated Ca²⁺ activity within dormant neural progenitors.
3. Ca²⁺-activated PAD enzymes link Ca²⁺ signalling to cell cycle progression by citrullinating intracellular targets in a Ca²⁺-dependent manner.
4. PAD inhibition suppresses medulloblastoma growth in preclinical laboratory models, revealing a conserved regulatory mechanism with translational potential.

## Main Text

Long considered a non-proliferative tissue after the cessation of embryonic neurogenesis, the vertebrate central nervous system (CNS) is now known to maintain a dynamic equilibrium between proliferation, quiescence, and senescence throughout life^1,2^. In the subgranular zone of the rodent dentate gyrus, for example, adult neurogenesis plays a crucial role in memory formation and cognitive plasticity^3^, and impaired adult neurogenesis has been observed in a range of neurodegenerative conditions^4^. Conversely, ectopic proliferation of cells with neural stem cell-like properties can give rise to primary brain and spinal cord tumours^5–8^, which represent some of the most aggressive and clinically intractable cancers. A deeper understanding of the mechanisms that control proliferation in the brain and spinal cord is therefore essential to our understanding of neural function and will underpin future efforts to treat proliferative disorders of the CNS.

In vertebrates, neurogenesis is typically driven by specialised ependymal radial glial cells (ERGs), which line the cerebrospinal fluid (CSF)-filled ventricles of the brain and spinal cord and serve as the primary pool of neural stem and progenitor cells in the CNS^9^. However, the neurogenic capacity of ERGs differs markedly across species^10^. For example, regenerative vertebrates such as teleost fish and salamanders show much higher rates of adult neurogenesis than most adult mammals and retain the ability to reactivate quiescent ERGs in response to injury throughout life^11,12^. Studies on regenerative systems therefore provide a tractable framework for dissecting the mechanisms that govern disease and repair and have been used to identify core regulatory mechanisms cells that are conserved across species^13–15^.

Here, we report the full trajectory from basic discovery to preclinical validation of a novel mode of proliferation control in the vertebrate CNS. Using high-resolution *in vivo* imaging of transparent larval zebrafish, we show that spinal cord injury induces acute changes in CSF circulation. These alterations reshape the local mechanical microenvironment, activating mechanoreceptors and triggering changes in intracellular Ca²⁺ dynamics that reawaken dormant ERGs. We identify the Ca²⁺-regulated peptidylarginine deiminase (PAD) family of enzymes as key intermediaries linking Ca²⁺ signalling to increased proliferation. Finally, we describe an analogous mechanism in human medulloblastoma cultures and demonstrate that PAD inhibitors effectively block tumour establishment and progression in a novel zebrafish xenograft model. Together, these findings uncover a hydrodynamic niche that regulates stem cell activity at the site of spinal cord injury in a regenerative vertebrate, highlighting the utility of regenerative systems for revealing mechanisms relevant to human pathologies of the CNS.

### Cilia-mediated CSF flow increases after injury

To uncover new mechanisms of quiescent ERG activation, we performed spinal cord injuries in larval zebrafish to trigger neurogenesis and examined signals that might couple injury to the subsequent reactivation of proliferation. Given the proximity of ERGs to the ventricular surface, and prior evidence that CSF contact influences proliferative behaviour in neurogenic niches across the vertebrate lineage and in cancers of the CNS^16–19^, we first asked whether CSF circulation is altered during regeneration and whether such changes could contribute toward ERG reactivation after injury.

Following partial spinal cord lesions (pSCL), which ablate the dorsal parenchyma but leave the central canal intact (Figure 1A to C), fluorescent microspheres injected into the hindbrain ventricle were rapidly observed in the circulating CSF of both uninjured and injured larvae (Movie S1), confirming the structural integrity of the central canal after pSCL. Quantitative analysis of particle trajectories in these samples revealed a significant increase in CSF flow within the central canal of injured larvae proximal to the lesion site compared to uninjured controls (Figure 1 D to K and Movie S2).

**Figure 1:**
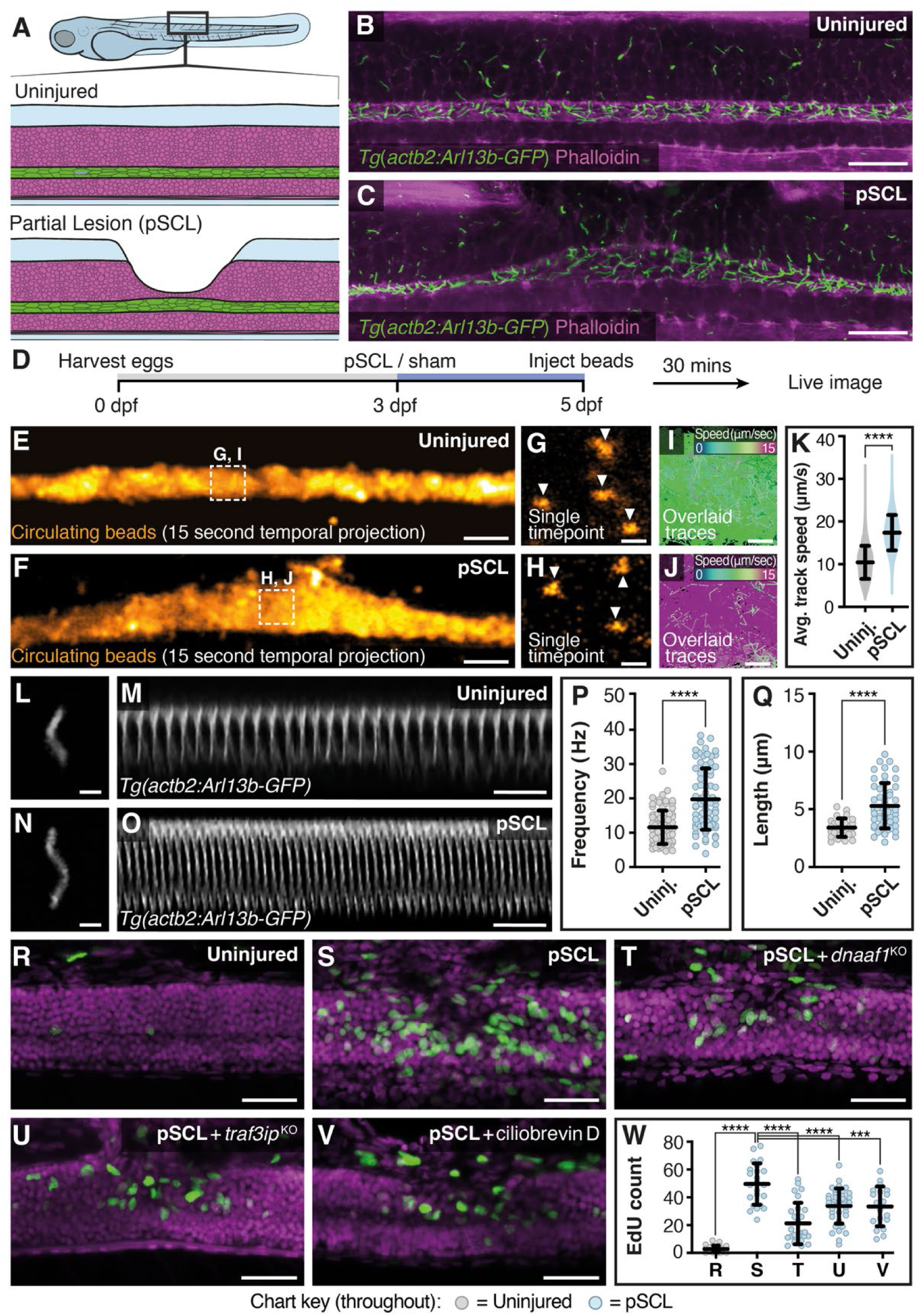
Cilia-driven CSF flow increases after injury and is required for regeneration. (A) Schematic of pSCL model. (B–C) Sagittal views of uninjured (B) and injured (C) larvae 1 h post-pSCL showing cilia marker *Tg(actb2:Arl13b-GFP)* (green) and F-actin (Phalloidin-555, magenta). Scale bars, 20 μm. (D) CSF flow quantification strategy. (E–F) Fifteen-second temporal projections of circulating beads in uninjured (E) and injured (F) larvae (see also Movie S1). Scale bars, 10 μm. (G–H) Single frames of circulating beads in uninjured (G) and injured (H) larvae. Scale bars, 2 μm. (I–J) Velocity-coded bead tracks from uninjured (I) and injured (J) larvae (see also Movie S2). Scale bars, 2 μm. (K) Average track velocity in uninjured and injured samples. (L–O) Single time-point images (L, N; scale bars, 2 μm) and 3 s kymographs (M, O; scale bars, 250 ms) of representative motile cilia in uninjured (L–M) and injured (N–O) larvae (see also Movie S3). (P) Motile cilia length at 48 hpi. (Q) Motile cilia beat frequency at 48 hpi. (R–V) EdU incorporation assays at 5 dpf (48 hpi) in uninjured (R), injured (S), injured + *dnaaf1* F0 knockout (T), injured + *traf3ip* F0 knockout (U), and injured + ciliobrevin D (V; 15 μM). (W) Quantification of R–V. In all figures, P < 0.05; P < 0.01; P < 0.001; P < 0.0001; ns, not significant. Middle lines indicate medians; top and bottom lines indicate standard deviation. Grey symbols denote uninjured samples and blue symbols denote pSCL.

CSF circulation is closely associated with the activity of flow-generating motile cilia, which beat asymmetrically to generate small-scale directional changes in CSF flow^20–24^. To determine if the observed increase in CSF flow after injury was caused by altered cilia dynamics, we performed partial spinal cord lesions on transgenic zebrafish ubiquitously expressing the fluorescent cilia marker *Arl13b-EGFP*^25^ (*Tg(actb2:Arl13b-EGFP)*) and utilised high speed, live fluorescence Airyscan imaging^26^ to record cilia activity in lesioned samples (Figure 1 L to O and Movie S3). Following injury, both the average length and the average beat frequency of motile ependymal cilia increased significantly in the vicinity of the lesion (Figure 1 P to Q), with the effect diminishing progressively with distance from the injury site (Figure S1).

Interestingly, the range of increased cilia activity broadly overlaps with the domain of ERGs that re-enter the cell cycle after injury^27^. To test whether increased CSF flow influences regenerative neurogenesis, we employed a series of pharmacogenetic interventions to disrupt motile cilia activity after injury and measured subsequent neural progenitor reactivation using 5-Ethynyl-2′-deoxyuridine (EdU) incorporation as a downstream readout of proliferation. Genetic perturbation of CSF circulation through CRISPR-Cas9 mediated F0 knockout^28,29^ of the intraflagellar transport protein *traf3ip1*^30^ or the axoneme assembly factor *dnaaf1*^31^, which are both essential for normal CSF flow^21^, resulted in reduced proliferation after injury (Figure 1 R to U). Similarly, treatment with the cytoplasmic dynein inhibitor Ciliobrevin D to disrupt motile cilia activity after injury^32,33^ significantly reduced proliferation (Figure 1 V). Collectively, these findings show that injury-associated changes in cilia-mediated CSF flow regulate ERG activation and proliferation in the regenerating zebrafish spinal cord.

### Locally acting inflammatory signals regulate cilia motility after injury

Molecules released from damaged or dying cells can act as inflammatory signals and are known to regulate tissue repair in the brain and spinal cord^27,34–38^. Inflammatory signals, including activation of the RNA-sensing receptor Toll-like receptor 3 (TLR3), are also known to influence cilia kinetics in ciliated epithelia^39^. Since double-stranded RNA (dsRNA) released from damaged cells activates TLR3 after spinal cord injury^27^, we next investigated whether TLR3 signalling regulates ependymal cilia motility in the lesioned spinal cord.

Treatment with a small molecule TLR3/dsRNA complex inhibitor (TLR3i) or RNAse A (to remove extracellular RNA species) significantly reduced cilia motility in the regenerating spinal cord (Figure 2 A). Conversely, injection of exogenous embryonic RNA extract into the CSF of uninjured larvae led to a significant increase in cilia beat frequency compared to vehicle-only injected controls (Figure 2 B to C). Increased cilia motility was not observed when the injected RNA extracts were pre-digested with RNase to disrupt secondary and tertiary RNA structures, which are essential for TLR3 activation^27,40^ (Figure 2 B to C). Together, these data demonstrate that RNA-induced activation of TLR3 signalling increases ependymal cilia motility.

**Figure 2:**
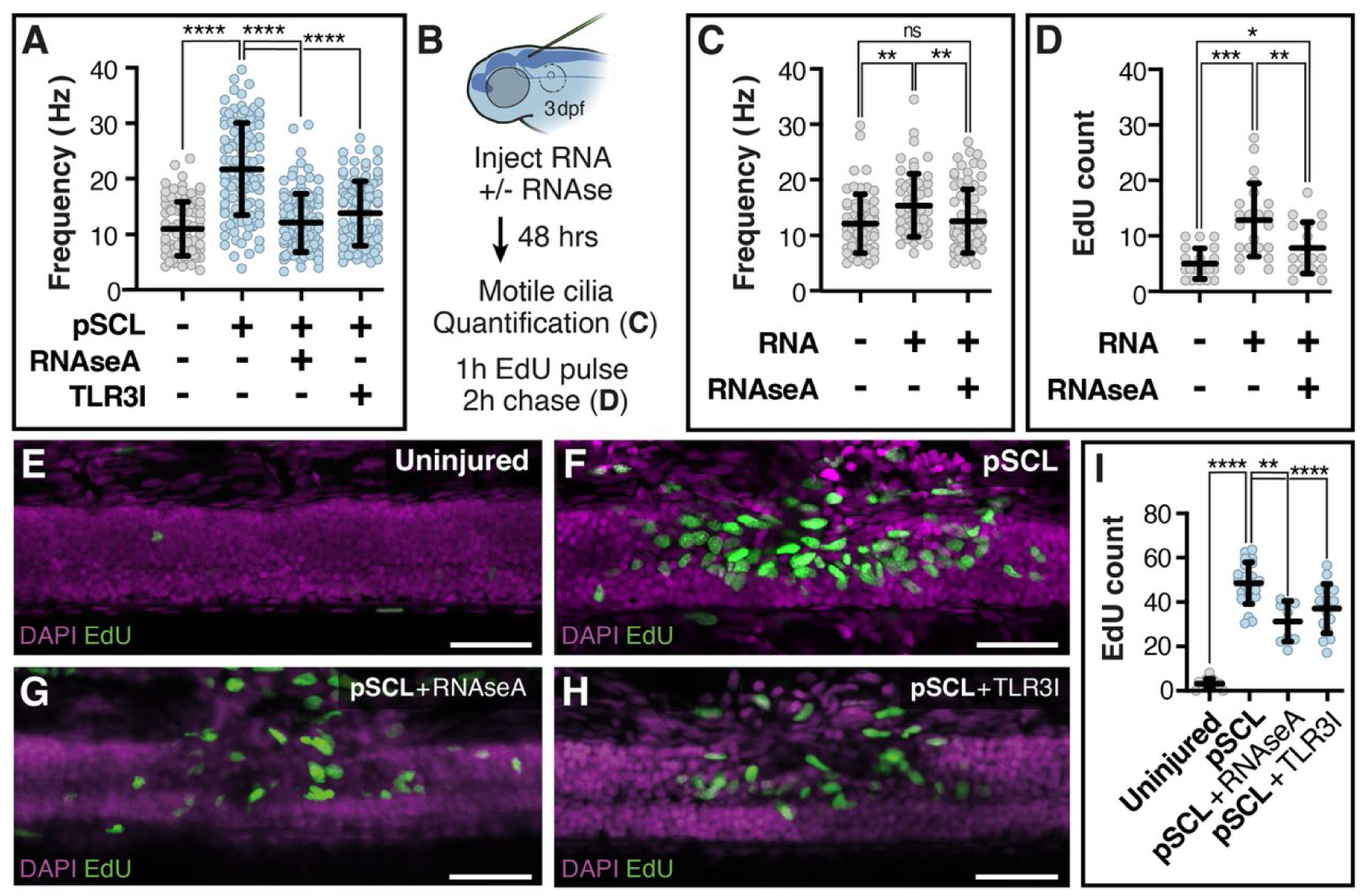
Wound associated inflammatory signals regulate cilia motility after injury. (A) Motile cilia beat frequency 48 h post-pSCL following treatment with RNase A (10 μg/ml) or the TLR3/dsRNA inhibitor TLR3i (2 μM). Images were taken proximal to the lesion site. (B) Experimental overview. (C) Motile cilia beat frequency in uninjured larvae following ventricular injection of total embryonic RNA (RNA) or co-injection with RNase A (10 μg/ml). Recordings were made 2 days after injection at 3 dpf. (D) EdU incorporation 48 h after ventricular injection of Ringer’s solution, total RNA, or RNA plus RNase A. (E–H) Representative EdU incorporation assays at 5 dpf in uninjured (E), injured (F), injured + TLR3i (G; 2 μM), or injured + RNase A (H; 10 μg/ml). Scale bars, 40 μm. (I) Quantification of E–H.

To demonstrate the role of damage-associated RNA signalling as an upstream regulator of CSF-circulation and ERG proliferation during regeneration, we first assessed whether increased cilia-mediated CSF flow was sufficient to initiate proliferation in the absence of injury. To do this, we injected RNA extract into the CSF of uninjured larvae (to activate TLR3 signalling in ependymal cells) and performed EdU incorporation assays to visualise subsequent ERG reactivation. Ventricular RNA injection significantly increased the number of proliferating cells in the spinal cord, and this effect was diminished upon co-injection of RNA and RNAse (Figure 2D).

To determine if RNA species released by damaged cells promote proliferation in the regenerating spinal cord, we treated injured larvae with TLR3i (to block TLR3 signalling) and RNAse A (to remove injury-associated extracellular RNAs). Both interventions showed significantly reduced proliferation after injury compared to untreated controls (Figure 2 E to I). Together, these data demonstrate that injury-associated inflammatory RNA species increase cilia motility and promote reactivation of quiescent ERGs near the injury site.

### Mechanosensitive neurons detect increased CSF flow after injury

In the vertebrate spinal cord, specialised mechanosensitive CSF-contacting neurons (CSF-cNs) detect alterations in CSF circulation and composition via specialised apical protrusions that extend into the lumen of the central canal^41–44^. To determine if CSF-cNs detect injury-associated changes in CSF flow, we performed population-level calcium imaging of larvae expressing the fluorescent Ca^2+^ sensor GCaMP6s^45^ under the control of the glutamate decarboxylase 1b promoter^46^ (*Tg(gad1b:Gal4/UAS:GCaMP6s)*). While this transgene labels all gamma-aminobutyric acid (GABA) producing neurons in the CNS, CSF-cNs can be specifically identified within this population by their characteristic apical protrusions, which colocalise with fluorescently labelled dextran tracers injected into the CSF (Figure 3 A to B and Movie S4).

**Figure 3:**
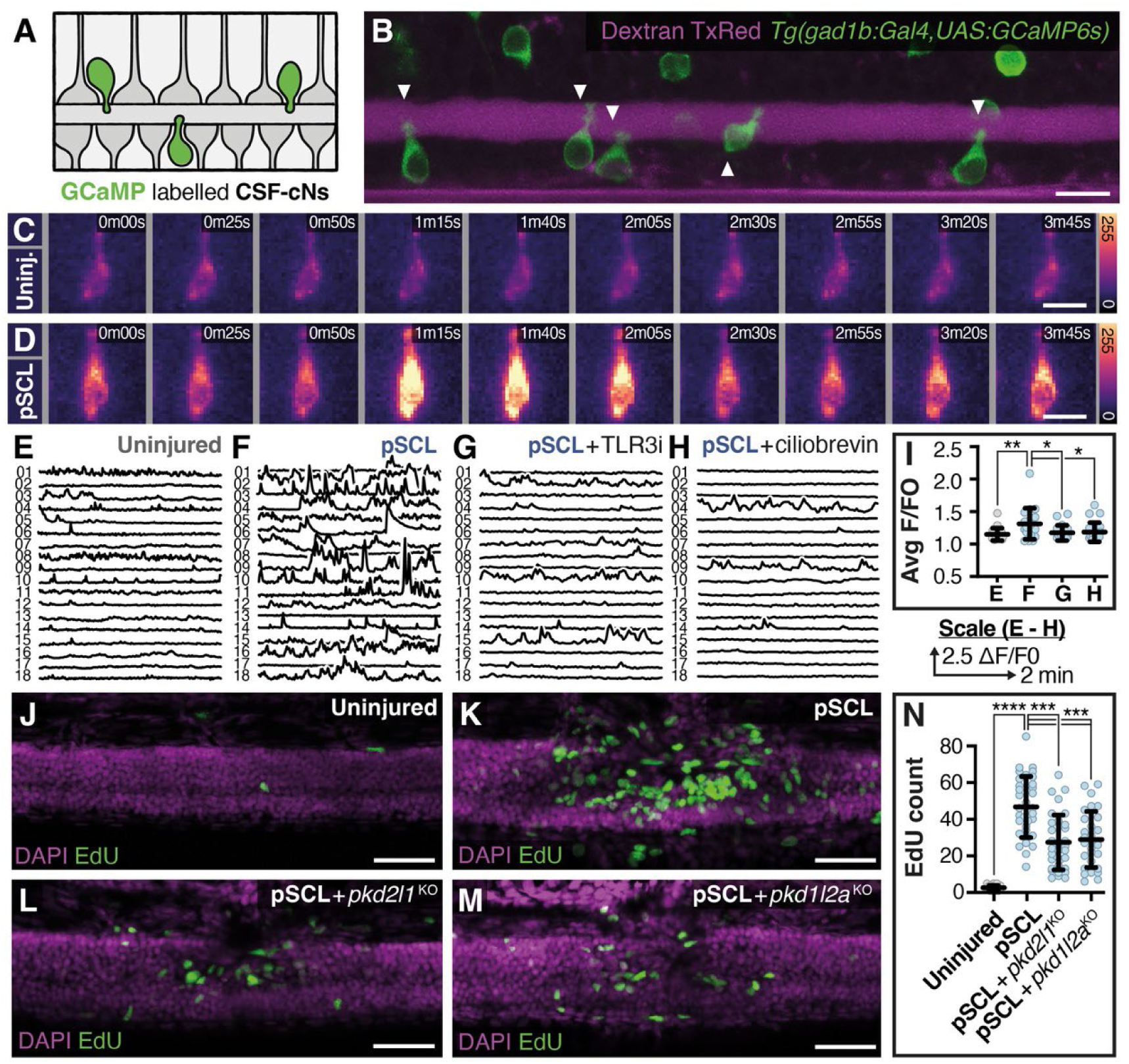
Mechanosensitive neurons detect elevated CSF flow during regeneration. (A) Schematic representation of cells labelled by *Tg(gad1b:Gal4/UAS:GCaMP6s* transgene. (B) Sagittal view of *Tg(gad1b:Gal4/UAS:GCaMP6s)* larva at 3 dpf injected with ∼1 nl of 70,000 MW Dextran TexasRed immediately before imaging to visualise cerebrospinal fluid. CSF-cNs (white arrowheads) are identified by their characteristic morphology and CSF-contacting apical processes (see also Movie S4). Scale bar, 10 μm. (C–D) Pseudo-coloured temporal series from high-speed time-lapse imaging of CSF-cNs expressing the Ca²⁺ sensor GCaMP6s in uninjured (C) and injured (D) larvae. Images were acquired at 5 dpf (2 dpi) following α-bungarotoxin paralysis. Scale bars, 5 μm. (E–H) Representative CSF-cN activity traces (normalised F/F₀) recorded at 5 dpf from uninjured (E), injured (F), injured + TLR3i (G; 2 μM), and injured + ciliobrevin D (H; 15 μM) larvae. Each trace represents a 10-min recording within 100 μm of the lesion site. (I) Quantification of average F/F₀ values from E–H. (J–M) Representative EdU incorporation assays (green) 48 h post-pSCL in uninjured (J), injured (K), and injured larvae following F₀ CRISPR knockout of *pkd1l2a* (L) or *pkd2l1* (M). Scale bars, 40 μm. (N) Quantification of J–M.

Limited spontaneous CSF-cN Ca^2+^ activity was observed in uninjured samples (Figure 3 C, E and I and Movie S5). In contrast, CSF-cN Ca^2+^ activity was significantly increased after injury (Figure 3 D, F and I and Movie S5). Elevated CSF-cN activity was not observed in injured samples following suppression of dsRNA-mediated TLR3 signalling with TLR3i (Figure 3 G and I), or inhibition of motile ciliogenesis with ciliobrevin D (Figure 3 H to I). Collectively, these data indicate that CSF-cNs respond to injury-associated changes in cilia-mediated CSF flow in the regenerating spinal cord.

Since cilia-mediated CSF circulation is required for ERG-reactivation after injury and CSF-cNs detect increased CSF-flow, we next investigated if CSF-cN activity is required for proliferation after injury. To do this, we performed CRISPR-Cas9 mediated F0 knockout of the mechanosensitive ion channel *pkd2l1*, a polycystin-2 (PC-2) family member which is expressed exclusively in CSF-CNs and is required for them to detect alterations in CSF circulation^42^. Following injury, *pkd2l1-*deficient larvae exhibited reduced proliferation compared to wild type controls (Figure 3 J to L and Figure 3 N). Similarly, F0 knockout of the polycystin-1 (PC-1) family protein *pkd1l2a*, which is co-expressed with *pkd2l1* in the zebrafish spinal cord and forms a likely PC-1/PC-2 mechanosensitive ion channel complex in CSF-cNs^47^, also resulted in significantly reduced proliferation after injury (Figure 3 M to N). Together, these data suggest that CSF-cNs detect injury-associated changes in CSF flow through the mechanosensitive ion channels *pkd1l2a* and *pkd2l1,* and that inhibition of this process limits the proliferation of ERGs in the regenerating spinal cord.

### Mechanosensitive neurons regulate ERG Ca^2+^ activity and proliferation after injury

CSF-cN activity is required for proliferation after injury, but proliferation is driven by the activation of quiescent ERGs. This suggests the existence of a neuron-to-glia signalling relay to link these two distinct populations within the spinal cord. Given that neuronally secreted factors promote gliogenesis in the early embryo by modulating calcium (Ca²⁺) levels in neurogenic zones^48–52^, and that Ca²⁺ is a well-established regulator of glial activation in both development and disease^53–57^, we focused on glial Ca²⁺ signalling as a potential activity-dependent mediator of ERG proliferation in the regenerating spinal cord.

We first determined if injury was associated with altered Ca^2+^ activity in ERGs by expressing GCaMP6s under the control of the *glial fibrillary acidic protein* (*gfap*) promoter (*Tg(gfap:Gal4/UAS:GCaMP6s)*), which labels radial glia in the zebrafish spinal cord^58^ (Figure S2 and Figure 4 A to B). Minimal ERG Ca^2+^ activity was detected in uninjured samples (Movie S6, Figure 4 G and K). However, after injury, strong waves of Ca^2+^ were observed throughout the lesion-proximal ERG domain (Movie S7, Figure 4, C to F, H and K). To determine if CSF-cN activity modulates the ERG Ca^2+^ response, we performed CRISPR-Cas9 mediated F0 knockout of *pkd2l1*, which abolishes flow sensing in CSF-cNs^42^. This significantly reduced injury-associated GCaMP6s activity (Movie S8, Figure 4 I and K), suggesting that CSF-cNs relay mechanical inputs to ERGs during regeneration.

**Figure 4:**
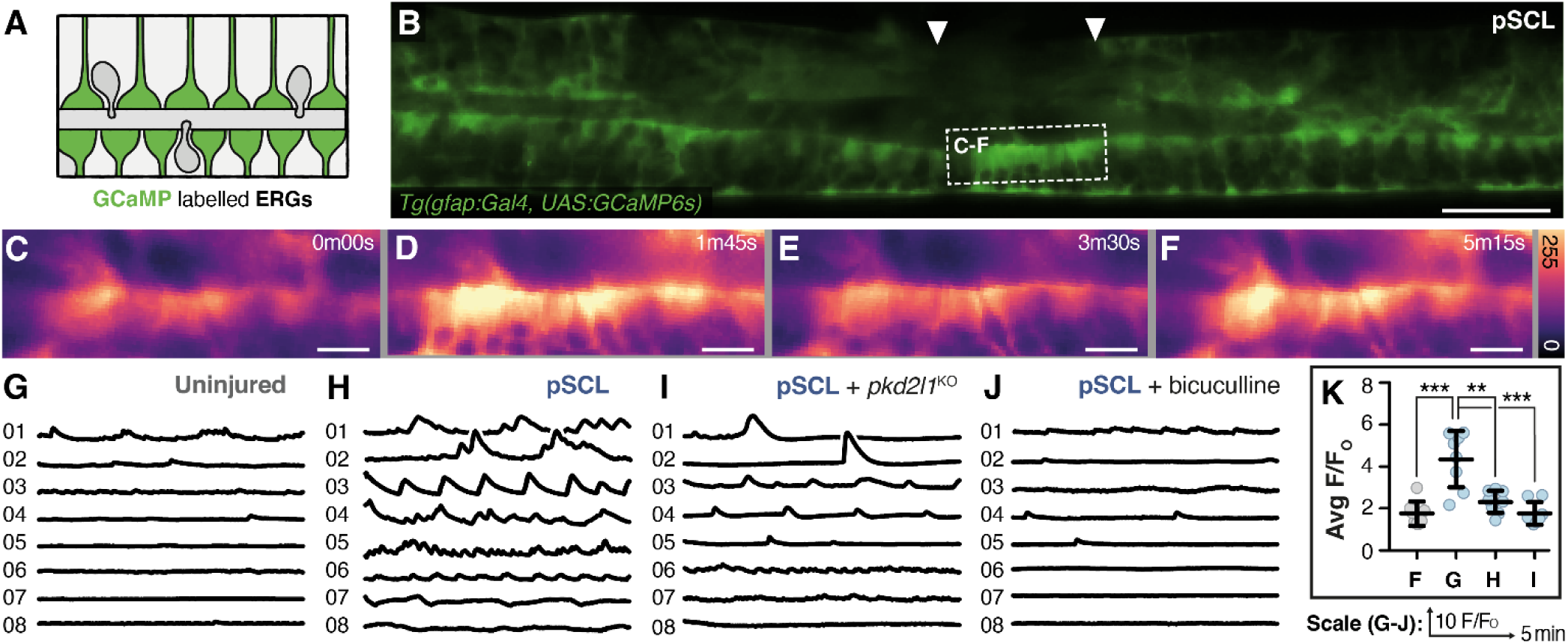
Mechanosensitive neurons regulate Ca²⁺ activity in ERGs. (A) Schematic representation of cells labelled by *Tg(gfap:Gal4/UAS:GCaMP6s* transgene. (B) Sagittal view of *Tg(gfap:Gal4/UAS:GCaMP6s)* larva at 48 hpl. White arrowheads indicate lesion boundaries, and the dashed box outlines the ependymal region shown in B–E. Scale bar, 40 μm. (C–F) Pseudo-coloured single-frame time points from high-speed time-lapse images of *Tg(gfap:Gal4/UAS:GCaMP6s)* larva at 48 hpl. Scale bars, 10 μm. (G–J) Fifteen-minute normalised *Tg(gfap:Gal4/UAS:GCaMP6s)* fluorescence traces from the lesion-proximal ERG region (∼100 μm upstream and downstream of the lesion site) at 5 dpf in uninjured (G), injured (H), injured following F₀ CRISPR knockout of *pkd2l1* (I), and injured following treatment with bicuculline (J; 25 μM). Each trace represents average ERG fluorescence across the gfap⁺ domain of a single larva. (K) Quantification of G–J.

Given that CSF-cNs produce and release GABA^59,60^ and that their activity is increased after injury, we next investigated whether GABA signalling is necessary for Ca^2+^ accumulation in ERGs within the regenerating spinal cord. Treatment with the GABA_A_R antagonist bicuculline reduced both injury-associated ERG Ca^2+^ activity (Movie. S9 and Figure 4 J to K) and proliferation (Figure S3) in the lesioned spinal cord, indicating that GABA is required for ERG reactivation.

Together, these observations support the existence of a transient hydrodynamic niche operating at the site of spinal cord injuries, where injury-associated dsRNA/TLR3 signalling increases motile cilia activity, stimulating local alterations in CSF flow and activating mechanosensitive ion channels on CSF-cNs at the wound site. In turn, CSF-cNs signal to ERGs in an activity-dependent manner, triggering Ca^2+^ transients that are essential for their proliferation.

### Ca^2+^ dependent citrullination promotes progenitor activity in the regenerating spinal cord

Ca²⁺ is an important signal in early wound healing, but how Ca²⁺ transients drive latent regenerative events such as the reactivation of quiescent progenitors remains unclear. To identify Ca²⁺-responsive factors involved in regeneration, we performed single-cell RNA sequencing after spinal cord injury. Activated ERGs were identified by *nestin* expression, which is strongly induced in proliferating neural progenitors following spinal cord injury (^27^ & Figure S4 A to B). This analysis revealed a subset of *nestin*-positive cells expressing the Ca²⁺-regulated enzyme *padi2* (Fig. S4 C to E).

The peptidylarginine deiminase (PAD) family of enzymes mediate protein citrullination, a post-translational modification that converts arginine residues into the non-coding amino acid citrulline^61,62^. Among the many known PAD substrates, histones have attracted particular attention, as their citrullination modulates chromatin accessibility^63–65^ and regulates the expression of several stem cell–associated genes^66–69^. To test whether citrullination is induced after injury in activated ERGs, we stained uninjured and injured samples with an antibody against citrullinated histone H4 (H4Cit3), a well-characterised PAD substrate previously used to detect citrullination in zebrafish^70^. Following spinal cord injury, we observed a strong and significant increase in histone citrullination at the wound site, with the most intense signal detected in ependymal cells adjacent to the lesion (Figure 5 A to B and E).

**Figure 5:**
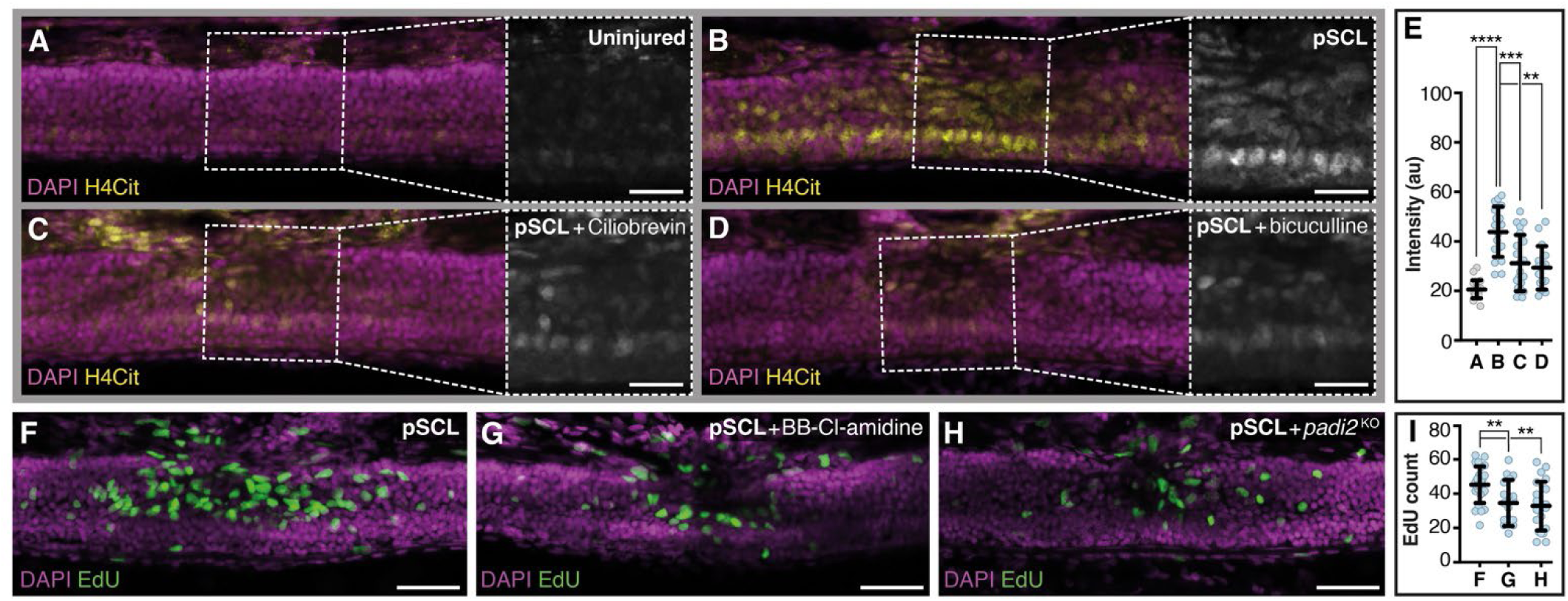
Ca²⁺-dependent histone citrullination is required for progenitor reactivation. (A–D) Representative images of citrullinated histone H4 immunoreactivity (H4Cit3, yellow) in uninjured (A), injured (B), injured + ciliobrevin D (C; 15 μM), and injured + bicuculline (D; 25 μM) larvae. Dashed boxes indicate regions magnified in insets. Scale bars, 20 μm. (E) Quantification of average H4Cit3 intensity across A–D. (F–H) Representative EdU incorporation assays (green) 48 hpi in control (F), injured + BB-Cl-amidine (G; 40 μM), and injured + *padi2* F0 knockout (H) larvae. Scale bars, 40 μm. (I) Quantification of F–H.

Since the activity of PAD enzymes is highly Ca²⁺-dependent^62^, we next asked whether the observed increase in citrullination arises from elevated Ca²⁺ levels after spinal cord injury. H4Cit3 immunostaining was performed on lesioned samples treated with Ciliobrevin D, to inhibit cilia motility (Figure 5 C), or bicuculline, to block CSF-cN–mediated Ca²⁺ accumulation in ERGs (Figure 5 D). Both compounds significantly reduced H4Cit3 immunoreactivity after injury (Figure 5 E), indicating that flow-associated changes in Ca²⁺ alter PAD activity during regeneration. To test whether PAD activity is required for progenitor activation and proliferation after injury, we inhibited PAD function with the PAD inhibitor BB-Cl-amidine or by CRISPR-Cas9 mediated F0 knockout of *padi2*, the only characterised PAD enzyme in zebrafish. Both interventions significantly reduced proliferation within the lesioned spinal cord (Figure 5 F to I), demonstrating that PAD-dependent citrullination promotes ERG activation during regeneration.

### Ca^2+^-dependent citrullination regulates medulloblastoma growth

ERGs in the zebrafish spinal cord share features with cerebellar stem and progenitor populations in zebrafish, as well as with cerebellar progenitors with regenerative potential in neonatal mice^27,71–73^. Intriguingly, medulloblastomas frequently arise from cerebellar progenitors and retain stem-like traits characteristic of neural stem and progenitor cells^74,75^. Based on these parallels, we next asked whether the Ca²⁺/PAD axis that drives ERG proliferation in the regenerating zebrafish spinal cord is also active in transformed cells.

To test this, we selected the DAOY cell line, a highly proliferative and invasive model of SHH-subgroup medulloblastoma^76^. Since PAD enzymes require Ca²⁺ to function, we first assessed whether spontaneous Ca²⁺ activity is present in DAOY cultures. Incubation with the cell-permeable Ca²⁺ indicator Fluo-4 AM revealed extensive, dynamic Ca²⁺ transients in confluent DAOY cultures. This activity was abolished upon treatment with the cell-permeable Ca²⁺ chelator BAPTA-AM, indicating that intracellular Ca²⁺ is elevated under basal conditions (Figure S5 and Movie S10).

Treatment of DAOY cultures with the PAD inhibitor BB-Cl-amidine led to a dramatic reduction in proliferation, as assessed by EdU incorporation (Figure 6A to E) and longitudinal cell counts (Figure 6F). Importantly, no increase in cell viability (Figure S6) or apoptosis (Figure S7) was detected upon PAD inhibition, indicating that Cl-amidine selectively impairs DAOY cell proliferation without inducing cytotoxicity. To assess whether PADs citrullinate histones in DAOY cells, we next quantified citrullinated histone levels by immunofluorescence. Strong nuclear staining was detected in untreated cells (Figure 6G and I) which was markedly reduced following BB-Cl-amidine treatment (Figure 6H to I), indicating active PAD-mediated histone citrullination. Together, these findings demonstrate that PAD enzymes citrullinate nuclear targets in medulloblastoma cells and that PAD activity is required for proliferation.

**Figure 6:**
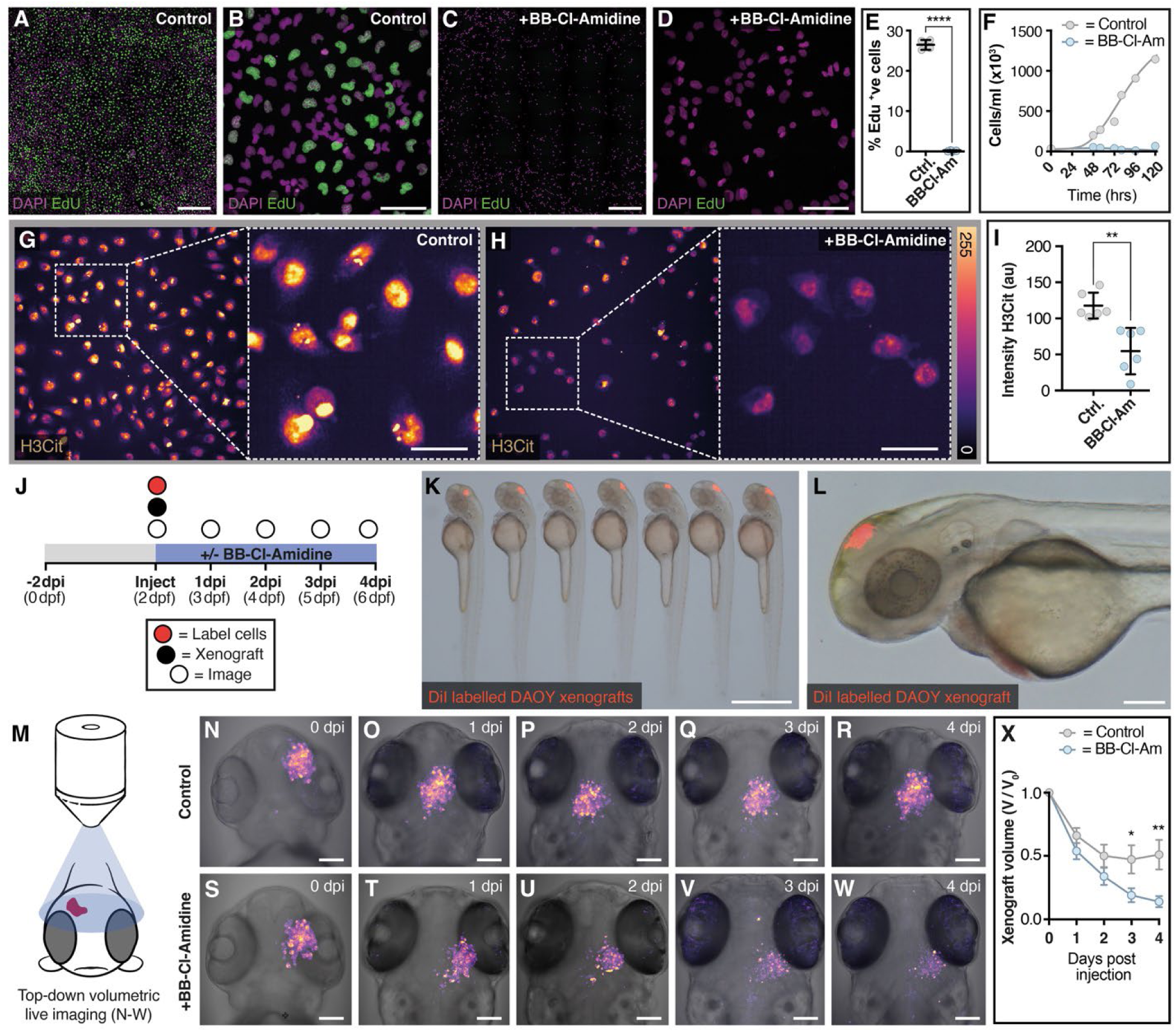
PAD-mediated citrullination promotes DAOY proliferation and supports establishment and expansion of larval xenografts. (A–D) Representative EdU incorporation assays (green) after 24 h treatment with DMSO (A, B) or BB-Cl-amidine (C, D; 6 μM). DAPI (magenta). Scale bars, 500 μm (A, B) and 100 μm (C, D). (E) Quantification of EdU incorporation in DAOY cells treated with BB-Cl-amidine or DMSO. (F) Growth-curve analysis of DAOY cell proliferation over 120 h following BB-Cl-amidine or DMSO treatment. (G–H) Representative maximum-intensity images of DAOY cells showing citrullinated histone H3 immunoreactivity after 24 h treatment with DMSO (G) or BB-Cl-amidine (H; 6 μM). Scale bars, 50 μm. (I) Quantification of average H4Cit3 intensity in (G) and (H). (J) Xenotransplantation experimental overview. (K) Side view of *DiI*-labelled DAOY cell pellets (red) transplanted into the midbrain parenchyma of 2 dpf embryos. Scale bar, 1 mm. (L) Magnified view of (K). Scale bar, 200 μm. (M) Imaging strategy for tracking xenograft volume over time. (N–W) Top-down maximum-intensity projections of fluorescently labelled DAOY xenografts from 0 to 4 dpi in untreated embryos (N–R) and embryos treated with BB-Cl-amidine (S–W; 40 μM). (X) Quantification of normalised tumour volume following DMSO or BB-Cl-amidine treatment.

To test whether PAD inhibition suppresses tumour growth *in vivo*, we developed a zebrafish neural xenograft model in which labelled cancer cells are transplanted into the midbrain parenchyma of 2-dpf larvae following ablation of microglia (the resident immune cells of the CNS^77^). This approach permits longitudinal imaging of xenografts to assess tumour growth dynamics and yielded highly reproducible transplants (Figure 6K), with labelled cells retained as spherical pellets at the injection site (Figure 6L). Serial volumetric imaging of transplanted xenografts revealed an initial reduction in tumour volume, followed by a period of stabilisation and expansion of the transplanted cells (Figure 6N to R). When the same experiments were conducted in the presence of BB-Cl-amidine, xenograft volume declined rapidly, with no evidence of stabilisation or regrowth (Figure 6S to X). Xenografts were reduced to minimal residual volume within four days of treatment (final volume <10% of baseline), demonstrating that PAD inhibition effectively suppresses the establishment and expansion of malignant medulloblastoma cells within the brain microenvironment.

## Discussion

Though considerable focus has been placed on the role of cerebrospinal fluid as a conduit for the transport of secreted factors within the developing and diseased CNS, the biomechanical implications of a circulating fluid system in direct contact with a neurogenic progenitor domain remain underexplored. Here, we show that CSF acts as a physical stimulus to establish a transient hydrodynamic niche at the site of spinal cord injury.

### Mechanosensory Regulation of Neural Progenitors

Our findings identify mechanosensitive CSF-contacting neurons (CSF-cNs) as crucial regulator of regenerative neurogenesis. This small and relatively understudied population of spinal sensory cells detects inflammation-associated alterations in CSF flow and transduces hydrodynamic cues to neighbouring glial populations in an activity-dependent manner. This signalling relay drives Ca²⁺ accumulation in ERGs, triggering neurogenesis and promoting repair. A similar reliance on Ca²⁺ signalling is seen in gap junction–coupled neural progenitors of the early neural tube, where coordinated Ca²⁺ transients propagate waves of developmental neurogenesis^78–80^. Intriguingly, *pkd2l1*-deficient mice display impaired neuronal regrowth and reduced motor recovery following injuries to the spinal cord^81^, raising the possibility that CSF-cNs also contribute toward mammalian CNS injury repair and response.

Though the broad roles of CSF-cNs in CNS physiology remain poorly defined, they are known to respond to a range of mechanical and chemical cues in the cerebrospinal fluid. For example, bacterial infection increases CSF-cN activity via activation of the orphan taste receptors Tas2r3a and Tas2r3b, but this does not lead to detectable Ca²⁺ accumulation in GFAP⁺ ERGs^43^. In contrast, physical activity^82^ and injury-induced alterations in CSF flow (this study) depolarise CSF-cNs via mechanosensitive Pkd2L1 channels and are associated with increased neurogenesis. This raises the intriguing possibility that chemical and mechanical inputs elicit distinct responses in CSF-cNs, with differential impacts on the status of neighbouring cells.

### RNA sensing initiates regenerative neurogenesis via TLR3-mediated modulation of CSF flow

Upstream of CSF-cN activation, we identify extracellular double-stranded RNA (dsRNA) released from damaged cells as an essential initiator of neural stem cell activity in the zebrafish spinal cord. We show that dsRNA acts through TLR3 to enhance ependymal cilia motility and locally increase CSF flow at the injury site, establishing a transient hydrodynamic niche that is permissive for ERG activation. This is consistent with the inflammatory modulation of motile cilia in the mammalian airway epithelium, where infection with RNA viruses triggers increased ciliary activity in a TLR3-dependent manner to promote airway clearance^39^. More broadly, our finding that extracellular dsRNA is required to initiate stem cell activation in the zebrafish spinal cord aligns with studies in mammalian systems, where wound-associated dsRNA activates TLR3 to promote skin regeneration and hair follicle neogenesis^83,84^, and where Tlr3-deficient mice exhibit delayed wound closure and impaired re-epithelialisation after wounding^85^.

We previously demonstrated that damage-associated dsRNA mediates the early, stem-cell–independent phase of zebrafish spinal cord repair by signalling through TLR3 to recruit primed precursor neurons, which rapidly restore circuit connectivity after injury^27,86^. Together with our current findings, these results establish TLR3 as a key regulator of both tissue remodelling and regenerative growth, and position RNA-sensing pathways as promising therapeutic targets in the CNS.

### Ca²⁺-dependent citrullination drives proliferation in regeneration and disease

Aberrant Ca²⁺ signalling is increasingly recognised as an important driver of tumour growth in the CNS, arising from diverse factors including synaptic input^87,88^, ion channel dysregulation^89,90^, and altered Ca²⁺ flux^91,92^. Recent work has also shown that GABAergic neuron-to-glia signalling elevates intracellular Ca²⁺ in transformed cells and promotes the progression of diffuse midline gliomas^93^. Although mechanistically distinct, the regulation of glioma progression by GABAergic interneurons bears many parallels with the activation of ERGs by GABAergic CSF-cNs during spinal cord regeneration.

How elevated Ca²⁺ drives proliferation of CNS tumours remains largely unclear. Here, we identify peptidylarginine deiminases (PADs) as Ca²⁺-dependent effectors of proliferation in both spinal cord regeneration and medulloblastoma. In zebrafish ERGs, injury activates PADs, which citrullinate intracellular targets such as histones at the lesion site. By studying medulloblastoma cells that share features with zebrafish ERGs, we show that PADs also citrullinate histones and promote proliferation in malignant cells. Importantly, PAD inhibitors suppress medulloblastoma growth effectively without inducing overt cytotoxicity and prevent tumour establishment and progression in zebrafish xenografts.

The role of PADs and citrullination in cancer is complex and context dependent. PAD activity has been implicated in the progression of breast, ovarian, prostate, and HPV-associated cervical cancers^94–97^, and PAD expression is associated with unfavourable clinical outcomes in cancers of the biliary duct^98^. Conversely, in colon cancer, PAD activation can have tumour-suppressive effects by inducing cell cycle arrest^99^. These differences likely reflect the diverse roles of citrullination as a post-translational modification, where functional outcomes depend not only on the identity of the modified protein but, in the case of histones, also on the specific genomic loci targeted. Elucidating the network of citrullinated proteins responsible for driving proliferation in different cell types and tissues will be an important focus of future research.

In summary, this study identifies a hydrodynamic signalling relay that links CSF flow dynamics to the reactivation of quiescent neural progenitors in the regenerating spinal cord. We show that increased CSF flow drives Ca²⁺ accumulation in ERGs via CSF-contacting neurons, leading to the activation of Ca²⁺-dependent PAD enzymes that citrullinate intracellular targets and initiate neurogenesis. A similar mechanism promotes proliferation in medulloblastoma cells, highlighting the potential of regenerative models for the identification of therapeutic candidates in the CNS.

## Acknowledgments

We thank many colleagues for support and advice during the course of this project; M. Montandon, J. Manneken and staff at Monash Micro Imaging for microscopy support; Monash FishCore staff for technical assistance and fish husbandry; B. Hogan and O. Baltaci for advice on flow tracking assays; B. Ciruna for the Tg(actb2:Arl13b-EGFP) construct; E. Scott for the Tg(gad1b:Gal4) and Tg(UAS:GCaMP6s) transgenic lines; M. Christophorou for advice on PADs and PAD related assays; C. Wyart for conceptual inputs and advice on *pkd2l1* knockouts.

## Author contributions

Conceptualisation: S.C. and J.K.

Methodology: S.C., T.W., A.D., C.F. and J.K.

Validation: S.C. and T.W.

Formal Analysis: S.C., T.W., A.D. and J.W.

Investigation: S.C., T.W. and J.W.

Resources: S.C., I.A. and A.D.

Data curation: S.C., T.W., A.D. and J.W.

Writing – original draft: S.C. and J.K.

Writing – review and editing: S.C., A.D., T.W. and J.K.

Visualization: S.C.

Supervision: G.L., I.A., S.C. and J.K.

Project administration: G.L., I.A., S.C. and J.K.

Funding acquisition: G.L., I.A., S.C. and J.K.

## Data and materials availability

Further information and requests relating to zebrafish resources and reagents should be directed to J.K.

## Supplementary Materials

Materials and Methods

Figs. S1 to S7

Movies S1 to S10

References 100-108

## Supplementary Materials

## Materials and Methods

### Zebrafish husbandry and fish lines

Fish were housed and bred within the Monash AquaCore Facility according to standard procedures^100^. All experiments were approved by the Monash Animal Services Animal Ethics committee, Monash University. The following fish strains used in this study were previously described: *Tg*(*actb2:Arl13b-EGFP)*^25^, *Tg(gad1b:Gal4)*^46^ and *Tg*(*UAS:GCaMP6s)*^101^.

Compound heterozygous *Tg(gad1b:Gal4/UAS:GCaMP6s)* larvae were generated by crossing *Tg(gad1b:Gal4)* and *Tg*(*UAS:GCaMP6s)* adults and screening progeny for green fluorescent signal within the expected *gad1b* expression domain using an Olympus MVX10 fluorescence stereomicroscope at 24 hpf.

### Generation of *gfap*:Gal4 line

The *Tg(gfap:Gal4)* line used in this study was generated by CRIMP mutagenesis^102^. The T2A-Gal4-VP16-synCoTC-4xnrUAS-mTagBFP2 targeting vector (Addgene plasmid ID: 199489) was selected to generate an in-frame Gal4 knock-in. A synthetic crRNA targeting the generic hBait site present in the CRIMP toolkit plasmids but absent from the zebrafish genome and a synthetic crRNA sequence targeting the first intron of the *gfap* locus were synthesised by IDT. For preparation of gRNAs, 2 μL of 100 μM crRNA was mixed with 1 μL of 100 μM tracrRNA (IDT) and annealed in a thermocycler at 95°C for 5 minutes, followed by a ramp down to 25°C at 5°C/min. An injection mixture containing 200 ng/μL gfap-targeting gRNA duplex, 100 ng/μL hBait-targeting gRNA duplex, 7.5 ng/μL targeting plasmid, 700 ng/μL Cas9 HiFi V3 protein (IDT, Cat. No. 1081061), 300 mM KCl, and 0.005% phenol red in sterile H₂O was incubated at 37°C for 1 h prior to injection, and 1–2 nL of the mixture was injected into 1-cell-stage wild-type embryos. Injected embryos were screened for mosaic mTagBFP2 expression in the expected *gfap* expression domain^58^, indicating successful integration at the *gfap* locus. Embryos displaying the highest level of mosaicism were reared to adulthood and outcrossed to *Tg*(*UAS:GCaMP6s)* to generate the compound heterozygous *Tg*(*gfap:Gal4/UAS:GCaMP6s)* larvae utilised in this study.

### Zebrafish CRISPR-Cas9 mediated F0 knockdown

F0 knockdown assays were performed according to a modified version of the protocol described in^28^. In brief, two synthetic crRNAs targeting distinct conserved coding regions within a given target gene were designed using the Alt-R CRISPR–Cas9 custom gRNA design tool (IDT). For preparation of gRNA duplexes, 2 μL of 100 μM crRNA was mixed with 1 μL of 100 μM tracrRNA (IDT) and annealed in a thermocycler at 95°C for 5 minutes, followed by a ramp down to 25°C at 5°C/min. The annealed guides were pooled and then an injection mixture containing 200 ng/μL genomic targeting gRNA duplex, 700 ng/μL Cas9 HiFi V3 protein and 0.005% phenol red in duplex buffer (IDT, 100 mM potassium acetate, 30 mM HEPES, pH 7.5) was prepared. 1-2 nL of injection mixture was injected into 1-cell-stage wild-type embryos and larvae were aged to experiment-appropriate time points (typically 3 dpf for spinal cord lesion assays).

### Zebrafish partial spinal cord lesion assay

The partial spinal cord lesion protocol used in this study was adapted from a previously described complete spinal cord lesion protocol^103^. Larvae were anaesthetised in 0.016% ethyl-m-aminobenzoate methanesulfonate (Tricaine; Sigma) at 3 days post fertilisation and embedded in 1% low-melting-point agarose on a 2% agarose-coated 90 mm Petri dish in a lateral orientation. A 0.38 mm tungsten wire needle (World Precision Instruments) was electrolytically sharpened and used to create mechanical lesions in the spinal cord at the level of the anal pore. Lesion width was approximately one myotome (∼100 μm) in diameter and care was taken to remove only dorsal tissue, ensuring the central canal remained intact. Lesions were performed under a backlit Nikon SMZ1500 stereomicroscope and samples with visible damage to the central canal were humanely euthanised in Tricaine and discarded. After lesion, larvae were gently released from the agarose using fine forceps and placed into fresh 1× Ringer’s medium (116 mM NaCl, 2.9 mM KCl, 1.8 mM CaCl₂, 5 mM HEPES in Milli-Q dH₂O, adjusted to pH 7.4) supplemented with 100 U/mL penicillin + 100 µg/mL streptomycin.

### Zebrafish ventricular injections

Larvae were anaesthetised in 0.016% ethyl-m-aminobenzoate methanesulfonate (Tricaine; Sigma) and embedded in 1% low-melting-point agarose on a 2% agar-coated 90 mm Petri dish, mounted in a lateral orientation. 1-2 nL of 125 µM α-bungarotoxin (2133, Tocris) was injected into the embryonic heart (as described in^73^). Samples were then gently released from the agarose using fine forceps, and loss of mobility was assessed to confirm efficient paralysis. Larvae were subsequently remounted on fresh 2% agar-coated dishes with the dorsal cranial surface facing upwards, and solutions were injected into the centre of the hindbrain ventricle.

For CSF flow tracking experiments, 1-2 nL of TetraSpeck microspheres were injected into the CSF of 5 dpf (2 dpi) larvae. Larvae were released from the agarose using fine forceps and remounted on 35 mm fluorodishes (World Precision Instruments) in a lateral orientation for immediate imaging (details below). This process was repeated for each larval sample individually to maintain a consistent interval between tracer particle injection and imaging of the circulating particles in the central canal.

For total RNA ventricular injections, RNA was isolated according to the protocol described in^104^. RNA concentration was determined with a NanoDrop One spectrophotometer (Thermo Fisher), adjusted to 500 ng/µL and injected into the centre of the hindbrain ventricle of 3 dpf larvae following the protocol described above. For RNA + RNase co-injections, 1 µL of RNase A (10 mg/mL, Thermo Fisher Scientific) was added to 20 µL of bulk RNA extract and incubated for 1 h at room temperature prior to injection.

### Drug treatments (zebrafish)

For zebrafish larval drug treatments, all compounds were diluted in 1× Ringer’s solution containing 0.5% DMSO, and the drug solutions were replaced daily. Control larvae received 1× Ringer’s solution with 0.5% DMSO only.

### Zebrafish 5-ethynyl-2’-deoxyuridine (EdU) labelling and detection

EdU labelling and detection in zebrafish larvae was performed as previously described^27^, with minor modifications. In brief, 5 dpf (2 dpi) larvae were pulsed with a solution of 50 μg/mL EdU and 5% DMSO in Ringer’s solution for 1 h, rinsed 5 times in 1× Ringer’s solution to remove residual EdU and DMSO and chased for 2 h in fresh 1× Ringer’s solution before fixation in 4% PFA in PBS overnight at 4°C. After permeabilisation in PBS-Tx (0.3% Triton X-100 in 1× PBS) for 2 h, EdU incorporation was detected by incubating samples in freshly prepared click-chemistry reaction buffer⁷⁵ containing 100 mM Tris (pH 8.5), 1 mM CuSO₄, 100 mM L-ascorbic acid, 5 µM Alexa Fluor 555 azide (Thermo Fisher Scientific, Cat. No. A20012), and 1 µg/mL 4′,6-diamidino-2-phenylindole dihydrochloride (DAPI) in 1× PBS. Samples were incubated overnight at 4°C, washed four times for 15 minutes each in PBS-Tx, and passed through a glycerol series (25%, 50%, 75% glycerol in PBS) prior to imaging.

### Zebrafish immunohistochemistry

For detection of histone citrullination in zebrafish, 5 dpf (2 dpi) larvae were fixed overnight at 4°C in 4% PFA in PBS and cleared using the CUBIC protocol^105^.

For detection of central canal morphology and integrity immediately after injury, 3 dpf (1 hpi) larvae were fixed in 4% PFA in PBS for 2 h at room temperature, washed thoroughly in PBS, and permeabilised in PBS-Tx for 30 min. Samples were incubated for 2 h at room temperature with Alexa Fluor 555 Phalloidin (Cell Signaling Technology, Cat. No. 8953) at the manufacturer’s recommended dilution and DAPI (1 µg/mL) diluted in PBS-Tx, then washed three times in PBS-Tx. Samples were passed through a graded glycerol series (25%, 50%, 75% in PBS), decapitated, and mounted in 75% glycerol for imaging.

### Tissue culture

DAOY medulloblastoma cells (ATCC, Cat. No. HTB-186) were maintained in Minimum Essential Medium Eagle (EMEM; Sigma-Aldrich, Cat. No. M0325) supplemented with 10% fetal bovine serum (FBS). Cells were cultured at 37°C in a humidified incubator with 5% CO₂ and passaged at a 1:5 ratio every 3–4 days using 0.25% trypsin–EDTA. Cell count and viability were assessed by trypan blue exclusion prior to seeding for experiments.

### Tissue culture drug treatments

For tissue culture drug treatments, DAOY cells were seeded and allowed to adhere overnight before treatment. The following day, the culture medium was aspirated and replaced with fresh complete medium containing BB-Cl-amidine or an equivalent volume of vehicle (0.1% DMSO).

### DAOY proliferation assay (growth curves)

DAOY medulloblastoma cells were seeded in 6-well tissue culture plates at a density of approximately 3×10⁴ cells per well on Day 0 to allow continuous proliferation without exceeding confluency by the final experimental time point. After 24 h, treatment wells were supplemented with BB-Cl-amidine in fresh complete medium, while control wells received an equivalent volume of vehicle (0.1% DMSO). After 24 h of treatment, one well from each condition was harvested, and viable cells were quantified by trypan blue exclusion using an automated cell counter (Countess™, Thermo Fisher Scientific). The remaining wells were replenished with fresh complete medium and incubated until subsequent collection time points. On each following day, one well per condition was trypsinised and counted as described above. Mean live cell counts were recorded for each time point (up to 120 h post-seeding), and growth curves were generated by plotting viable cell numbers against time to assess proliferation dynamics.

### Fluorescence-activated cell sorting (FACS)

DAOY medulloblastoma cells were seeded in 6-well plates at a density of 1.5–2.0×10⁵ cells per well, cultured overnight, and treated with BB-Cl-amidine or 0.1% DMSO for 24 h. Staurosporine (2 µM, 4 h) was used as a positive control to induce apoptosis, and Triton X-100 (0.1%, 15 min) was used as a positive control for cell viability assays to disrupt membrane integrity. Cells were harvested, pelleted at 300 × *g* for 5 min, and stained with FITC–Annexin V and propidium iodide (PI) according to the manufacturer’s protocol (Thermo Fisher Scientific, Cat. No. V13241A/B). Hoechst 33342 (2 µg/mL) was included to label nuclei. For membrane integrity assays, only Hoechst and PI were used. Controls included unstained and single-stained samples (Hoechst, Annexin V, or PI). Staining was performed for 15–20 min at room temperature in the dark, followed by two PBS washes.

Data were acquired and analysed using a BD LSRFortessa™ X-20 flow cytometer and FlowJo v10 software (BD Biosciences). For each sample, a minimum of 30,000 events was collected.

For viability assays (Hoechst + PI), a three-step gating strategy was applied: (1) forward scatter area (FSC-A) versus side scatter area (SSC-A) plots were used to exclude debris and identify the main cell population; (2) singlets were gated based on forward scatter height (FSC-H) versus area (FSC-A) to remove aggregates; and (3) Hoechst versus PI plots were used to distinguish membrane-compromised cells (Hoechst⁺/PI⁺) from viable nuclei (Hoechst⁺/PI⁻).

For apoptosis assays (Annexin V + PI + Hoechst), a four-step gating strategy was used: (1) debris exclusion by FSC-A versus SSC-A; (2) singlet discrimination by FSC-H versus FSC-A; (3) selection of nucleated cells based on Hoechst fluorescence; and (4) Annexin V versus PI plots to classify viable (Annexin V⁻/PI⁻), early apoptotic (Annexin V⁺/PI⁻), and late apoptotic or necrotic (Annexin V⁺/PI⁺) populations.

Fluorescence thresholds were defined using unstained and single-stained controls for each fluorophore. All conditions were assayed in triplicate.

### DAOY larval xenografts

DAOY medulloblastoma cells were seeded in T25 flasks at a density of 1–1.5×10⁶ cells to reach approximately 70% confluency within 24 h. Following incubation, the culture medium was aspirated and cells were gently washed with PBS for 30 s to 1 min with gentle swirling to remove residual serum. Cells were then labelled with Vybrant™ DiI cell-labelling solution (Thermo Fisher Scientific, Cat. No. V22885) by adding 10 µL of 2 µM dye solution to 5 mL of serum-free complete growth medium and incubating for 30 min at 37°C. After incubation, the dye-containing medium was aspirated and replaced with 5 mL of fresh complete growth medium. Cells were further incubated for 10 min at 37°C to allow dye equilibration and then washed three times with PBS to remove excess dye. Labelled cells were harvested using 0.25% trypsin–EDTA, and cell count and viability were assessed by trypan blue exclusion. The resulting cell pellet was resuspended in 30–50 µL of polyvinylpyrrolidone (PVP; Merck Life Science, Cat. No. PVP40) in a 1.5 mL microcentrifuge tube, with the volume adjusted according to pellet size. The injection mixture was protected from light and maintained at 37°C until use for xenograft injections.

Since transient downregulation of immune cells has previously been shown to enhance zebrafish xenograft engraftment^106^, we performed CRISPR–Cas9-assisted F0 knockout of *interferon regulatory factor 8* (*irf8*) to suppress microglia^77^ prior to xenotransplantation. Following knockdown, embryos were cleaned daily. At 48 hpf, *irf8*-deficient larvae were anaesthetised in tricaine, embedded in 1% low-melting-point agarose, and mounted on 2% agar-coated 90 mm Petri dishes in an upright orientation with the dorsal surface facing upward to facilitate injection into the midbrain parenchyma. DiI-labelled DAOY cells were loaded into a pulled glass capillary needle, and a pellet containing approximately 100–150 cells was injected into the right hemisphere, with manual injection volumes determined by visual assessment of pellet size. Following injection, larvae were gently released from the agarose using fine forceps and transferred to individual wells of a 12-well plate containing 1× Ringer’s solution supplemented with 6 µM BB-Cl-amidine or 0.5% DMSO (vehicle control) and maintained at 35°C. After xenotransplantation, larvae were immediately mounted individually for live volumetric imaging and then returned to the 12-well plate after imaging. Imaging was repeated at 24-h intervals for four consecutive days (see Imaging section for details).

### DAOY 5-ethynyl-2’-deoxyuridine (EdU) labelling and detection

For EdU detection and labelling, DAOY cells were seeded at a density of 2×10⁴ cells per well in tissue culture-treated 96-well optical-bottom black microplates and incubated overnight to allow cell attachment. Cells were then treated with either 0.1% DMSO (vehicle control) or 6 µM BB-Cl-amidine for 24 h. Following treatment, the culture medium was replaced with fresh medium containing 20 µM EdU, and cells were incubated for an additional 2 h at 37°C. After EdU incorporation, the medium was aspirated and cells were fixed immediately in 4% PFA for 15 min at room temperature, washed, and permeabilised with 0.1% Triton X-100 for 15 min. EdU signal was detected by incubating cells with freshly prepared click-chemistry reaction buffer (as described above) for 1 h at room temperature. After incubation, cells were washed three times with PBS and imaged.

### DAOY immunohistochemistry and live dye staining

For detection of histone citrullination in medulloblastoma cultures, DAOY cells were seeded and incubated overnight in black, tissue culture-treated 96-well optical-bottom black microplates at a density of 3×10⁴ cells per well in the presence of 6 µM BB-Cl-amidine or vehicle control (0.1% DMSO) for 24 h (four replicate wells per condition). Following treatment, cells were fixed in 4% PFA for 15 min at room temperature, permeabilised in 0.1% Triton X-100 for 15 min, blocked for 1 h in 2% BSA in PBS, and stained with anti-H3Cit antibody overnight at 4°C. Samples were washed three times in PBS and incubated for 2 h at room temperature with an Alexa 555-conjugated secondary antibody and DAPI (1 µg/mL) in PBS. Samples were washed three times in PBS and imaged.

To visualise Ca²⁺ dynamics in DAOY cultures, cells were seeded in 35-mm glass-bottom fluorodishes at a density of 3×10⁵ cells per dish and incubated overnight to allow adhesion to the glass surface. The following day, cells were incubated with the cell-permeable calcium indicator Fluo-4 AM (5 µM; Thermo Fisher, Cat. No. F14201) in standard culture medium for 10–15 min at 37°C. After incubation, cells were washed three times in fresh medium before imaging. Baseline fluorescence was recorded for 6 min, after which 100 µL of 100 µM BAPTA-AM was added per dish to reach a final concentration of 10 µM and establish a calcium-free baseline.

### Imaging

Fixed and immunostained zebrafish samples were imaged using a Leica Stellaris 5 inverted confocal microscope with a 20×/0.75 NA multi-immersion objective. Stacks were acquired with a 1.5 µm step size, spanning the entire transverse aspect of the spinal cord.

For bead-based flow-tracking experiments, α-bungarotoxin-paralysed larvae were mounted in 1% low-melt agarose on 35 mm fluorodishes and imaged at 560/580 nm (ex/em) using a Leica Stellaris 5 inverted confocal microscope with a 20×/0.75 NA multi-immersion objective immediately after microsphere injection (see above). To enhance acquisition frame rates, a single Z-plane was acquired, allowing images to be collected at 30 fps (33.33 ms per frame).

High-temporal resolution imaging of motile cilia was performed on a Zeiss LSM 980 upright confocal microscope equipped with an Airyscan 2 imaging module and a 20×/1.0 NA water-immersion objective. Images were captured in Fast Airyscan mode (Fast-AS) with default Airyscan filtering, enabling an acquisition rate of 200 fps (4.98 ms per frame).

Whole spinal cord GCaMP6s imaging (using *gad1b-* and *gfap-*driven Gal4 drivers) was conducted using a Zeiss LSM 980 upright confocal microscope with a 20×/1.0 NA water-immersion objective. To balance acquisition speed with optical section thickness, the pinhole was set to 5 Airy units. To ensure a sufficient transverse aspect of the spinal cord was captured, a Z-stack of five slices, spaced 7.5 µm apart, was centred on the central canal and continuously acquired over a 10-minute period, with subsequent analysis performed on maximum intensity projections. This setup yielded an acquisition rate of 0.4 fps (2.5 seconds per frame).

Longitudinal volumetric imaging of DAOY xenografts was performed using a Zeiss LSM 980 upright confocal microscope equipped with a 20×/1.0 NA water-immersion objective. Live larvae were anaesthetised in 0.016% tricaine and embedded in an upright (“head-up”) orientation for imaging. Z-stacks spanning the full depth of the injected cell mass were acquired. After imaging, samples were gently demounted and returned to individually housed wells of a 12-well plate containing 1× Ringer’s solution with or without drug treatment. The same imaging protocol was repeated daily until the final experimental time point at 4 dpi (6 dpf).

Time-lapse imaging of Fluo-4–loaded DAOY cells was performed on a Leica Stellaris 5 inverted confocal microscope equipped with a 20×/0.75 NA multi-immersion objective. Single z-planes were acquired at 1 s intervals. BAPTA-AM was added mid-acquisition by gentle pipetting.

Imaging of EdU-labelled and H3Cit-stained DAOY cultures was performed using the Opera Phenix™ high-content confocal microscope (PerkinElmer). Images were acquired using a 63× objective in confocal mode with laser-based autofocus. Forty-nine non-overlapping fields were captured per well, and imaging parameters were kept constant across all conditions.

### Image analysis

Microscopy images were processed in FIJI and Imaris 10.2 (Bitplane). To quantify motile cilia beat frequency, time-lapse datasets were imported into FIJI, and a straight line was drawn from the base to the tip of each cilium. Kymographs were generated using the Reslice function, and beat frequency was quantified by segmenting the kymographs with the Threshold command. Individual beats (visible as horizontal white bands across the image) were counted using the Analyse Particles command, with each band corresponding to a complete ciliary beat cycle.

To quantify H4cit3 intensity, maximum intensity Z-projections were generated in FIJI, and images were rotated to ensure the spinal cord was positioned in a horizontal alignment across the image. The image was then cropped to a width of 300 µm to define the lesion-proximal portion of the spinal cord used in subsequent analyses. Regions of interest (ROIs) were drawn around the spinal cord and a cell-free area of the notochord (serving as a background reference to account for sample-to-sample variation in background fluorescence) with the DAPI counterstain used to define the boundaries of each ROI. Average intensity values were measured for both the spinal cord and background ROIs, with the background intensity subtracted from the spinal cord intensity to calculate the normalised fluorescence values reported in this study.

For EdU counts, maximum intensity Z-projections were generated in FIJI, rotated to ensure the spinal cord was positioned in a horizontal alignment and cropped to a width of 300 µm as described above. EdU+ cells within the spinal cord (defined according to the DAPI counterstain) were manually counted using the native Cell Counter plugin.

To generate *Tg(gad1b:Gal4/UAS:GCaMP6s)* intensity traces, maximum intensity Z-projections for all time points were generated in FIJI. If present, motion artefacts caused by slow drift of the sample were corrected using the Fast4DReg plugin^107^. ROIs were drawn around individual CSF-cNs (identified by their proximity to the central canal and characteristic morphology) and average intensity values were acquired for each time point using the ROI multi-measure tool. F0 was defined as the minimum value within each dataset and F/F0 was calculated by dividing the average intensity value measured at each time point by F0.

To generate *Tg(gfap:Gal4/UAS:GCaMP6s)* intensity traces, maximum intensity Z-projections were generated for all time points, and motion artefacts were corrected as necessary using Fast4DReg^77^. A single ROI was drawn around the spinal cord and average intensity values were acquired for the entire tissue. To account for fluorophore bleaching during extended acquisition periods (>15 minutes), fluorescence was normalized by identifying local minima and correcting for baseline decay using polynomial regression. F0 was defined as the minimum value within each dataset and F/F0 was calculated by dividing the average intensity value measured at each time point by F0.

Quantification of EdU in DAOY cells was performed using the in-built Harmony analysis software (v2.1) of the Opera Phenix™ high-content confocal microscope. EdU-positive nuclei were identified using fluorescence intensity thresholding, and the percentage of EdU-positive cells was calculated relative to the total number of DAPI-stained nuclei.

H3Cit quantification in DAOY cultures was performed in Fiji. Nuclear masks were generated from the DAPI channel and used to measure mean H3Cit fluorescence intensity across each nucleus.

To generate Fluo-4 intensity traces in DAOY cultures, ROIs were manually drawn around individual cells, and mean fluorescence intensity values were measured for each time point using the multi-measure tool in Fiji. F0 was defined as the minimum intensity value within each dataset, and F/F0 was calculated by dividing the fluorescence intensity at each time point by F0.

Xenograft volumes were calculated using the in-built Surfaces tool in Imaris (Bitplane, Oxford Instruments) with automated thresholding. Volumes were normalised to the initial xenograft volume (1 h post-injection) for each larva tracked throughout the experiment.

### Single-cell sequencing: spinal cord injuries and tissue collection

Double-transgenic adult zebrafish (*Tg^KI^(glula-T2A-EGFP);Tg(-8.7elavl3:mCherry-T2A-CreERT2)*)^27^ were deeply anaesthetised in 0.033% Tricaine methanesulfonate. The vertebral column was exposed by a longitudinal incision on the dorsum at the level of the anal pore, and the spinal cord completely transected using fine forceps with care taken to leave the underlying vasculature intact. Zebrafish were recovered from anaesthesia in system water, and monitored daily until their experimental endpoint. At 1- and 3-days post-lesion (dpl), zebrafish were humanely euthanised in an ice-water slurry, and the spinal cord region proximal to the lesion site was dissected in cold 1x PBS. Spinal cords from uninjured (control) animals were dissected as above at the same anatomical level. 8 spinal cord regions were pooled for dissociation per group.

Tissue was rinsed in ice-cold 1x HBSS, then incubated for 15 mins at 37 °C in 1 mg/mL collagenase II in 0.1 M Tris-Cl pH 7.5 while shaking at 300 rpm. Samples were gentle triturated by repeat pipetting through a wide-bore 1000 µL pipette tip every 5 mins during the dissociation. Cells were pelleted by centrifugation at 2000 rpm for 5 mins at 4 °C, then resuspended in 1 mL HBSS + 2% fetal calf serum (FCS), pelleted again, and resuspended in 500 µL HBSS + 2% FCS. Samples were passed through a 35 µm cell strainer; cell number and viability were quantified using Trypan Blue staining of a 5 µL cell aliquot, and the final viable cell concentration adjusted to 1 x 10^6^ cells/mL.

### Single-cell sequencing: library preparation, and sequencing

100 bp paired-end libraries were prepared using Chromium Single Cell 3’ V3.1 chemistry (10X Genomics), and sequenced on an MGISEQ2000RS platform (MGITech) using MGIEasy V3 chemistry by the Monash Genomics and Bioinformatics Platform (MGBP, formerly Micromon). Each sample produced between ∼140-195 million reads. The sequencing output was processed using the CellRanger pipeline (v6.1.1) to generate feature-barcode matrices. Base calls were demultiplexed into FASTQ reads, which were aligned using cellranger count to the zebrafish GRCz11 genome containing a custom chromosome representing GSG-T2A-EGFP and mCherry-T2A-CreERT2 ORFs corresponding to the two transgenes. Only exonic reads were included during alignment. Filtered feature-by-cell matrices output from CellRanger’s cellranger count function were used for subsequent analyses.

### Single-cell sequencing: data processing and analysis

Data were integrated, quality-controlled and analysed using Seurat (v5.3.0)^108^ in R (v4.5.1). Seurat objects were created from each sample’s filtered CellRanger outputs, and initially filtered to remove genes expressed in fewer than 2 cells, and to remove cells expressing fewer than 100 genes. Further quality control was performed by subsetting each object to exclude obvious doublets based on number of expressed features and RNA molecules (determined on a per-sample basis). Cells where reads derived from mitochondrial genes exceeded 20% of that cell’s total transcriptome were excluded. This yielded 380, 559 and 1,049 high-quality cells for control, 1 dpl, and 3 dpl groups, respectively. Each sample aggregated into a single merged object which was then log-normalised, and the top 2000 variable features were identified using the ‘vst’ method. The data were then scaled, and PCA was performed using the top 20 principal components (PCs) as determined by an Elbow plot. These 20 PCs were used for nearest-neighbour graph construction and dimensionality reduction using the UMAP method with parameters ‘spread = 0.2’, ‘min.dist = 0.1’ and ‘learning.rate = 1). Clustering was performed using the Louvain algorithm, and the clustree package was used to identify the optimal cluster resolution (0.4) to clearly resolve distinct cell populations. Cluster marker genes were identified using a Wilcoxon rank sum test via the ‘FindAllMarkers’ function with parameters ‘min.pct = 0.15’, ‘min.diff.pct = 0.15’ and ‘only.pos = T’. These cluster markers were used to assign biological identity for all clusters. Basic dimensionality reduction visualisation was performed using Seurat functionalities as well as the SPRING web interface.

### Manuscript and figure preparation

Figures were compiled using Adobe Photoshop and Adobe Illustrator. Fluorescent and confocal microscopy images were adjusted globally for brightness and contrast using FIJI and then flattened into RGB images and exported as TIFFs.

### Statistics

Larvae for experimental treatment groups were randomly assigned and sample sizes were not predetermined with statistical methods. All experiments included at least three independent biological replicates, and the number of samples used is provided in the figure legends. Plots show individual datapoints, overlaid bar charts display mean value (horizontal central line) and standard deviation (upper and lower error bars). Statistical analysis was performed in Prism (GraphPad Software). Data was analysed using Student’s unpaired two-tailed *t*-test with pairwise comparisons denoted by brackets on charts. For all graphs, ns = not significant, * < 0.05, ** < 0.01, *** < 0.001 and **** <0.0001.

**Figure S1:**
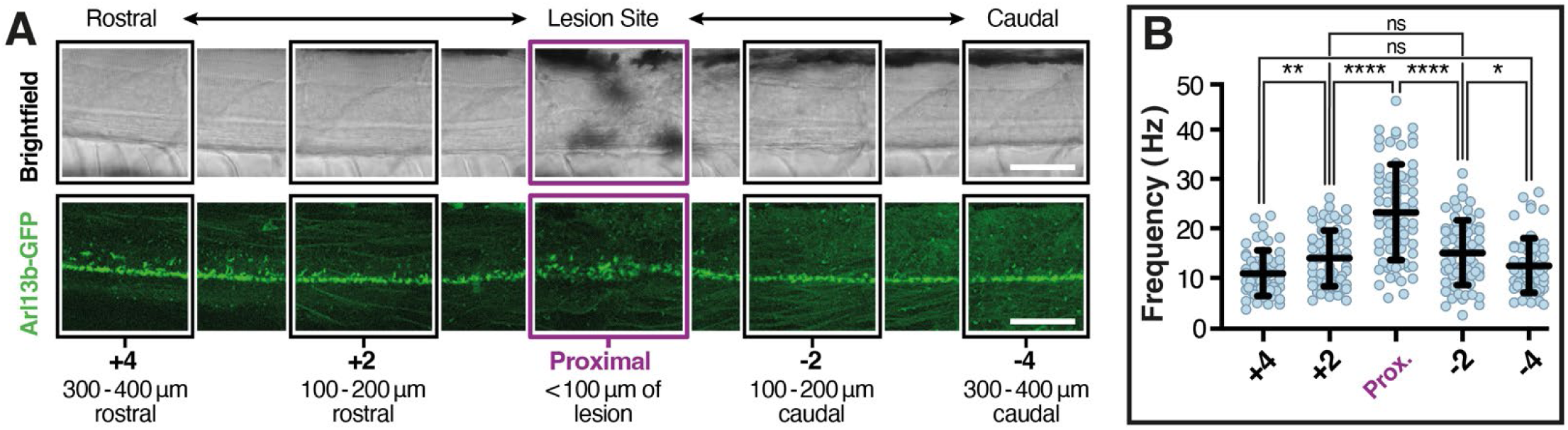
Increased cilia motility is localised to the lesion site. (A) Sagittal view of *Tg(actb2:Arl13b-GFP)* larva at 2 dpi. The lesion site is outlined in magenta, and distal segments are highlighted in black. Numerical labels (+4, +2, –2, –4) indicate the number of segments rostral (+) or caudal (–) to the lesion. These bins were used to map changes in cilia beat frequency relative to the lesion. CC, central canal. Scale bar, 50 μm. (B) Quantification of motile cilia beat frequency 48 h post-pSCL within the 100 μm bins described in (A).

**Figure S2:**
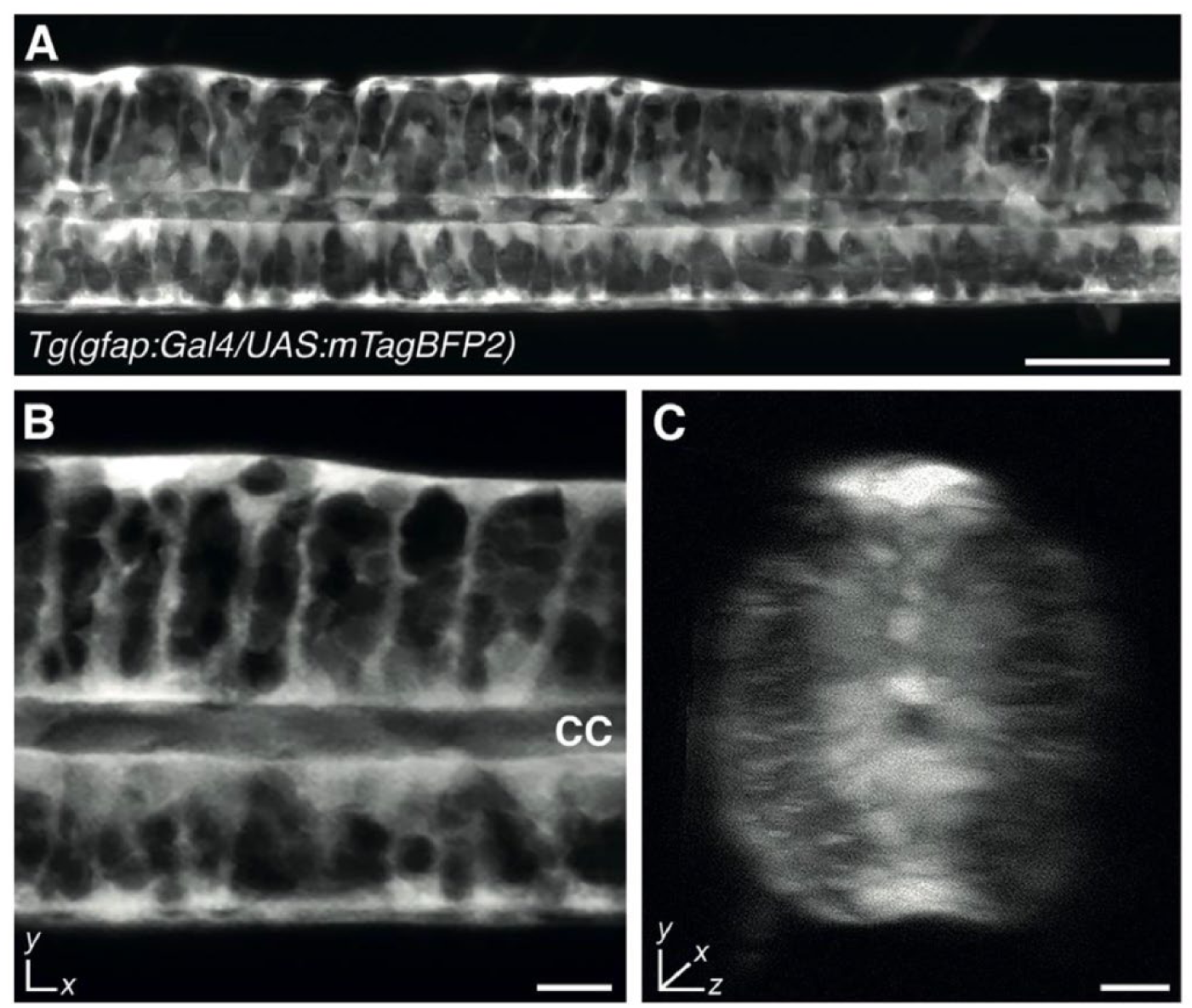
Endogenous Tg(gfap:Gal4) knock-in reporter labels spinal cord ERGs. (A) Sagittal view of an uninjured *Tg(gfap:Gal4/UAS:mTagBFP2)* larva. Image is a maximum-intensity Z-projection spanning the full depth of the spinal cord. Scale bar, 40 μm. (B) Single Z-slice showing *Tg(gfap:Gal4/UAS:mTagBFP2)*-labelled glia with characteristic radial morphology. CC, central canal. Scale bar, 10 μm. (C) Orthogonal projection of the dataset shown in (A). Scale bar, 10 μm.

**Figure S3.**
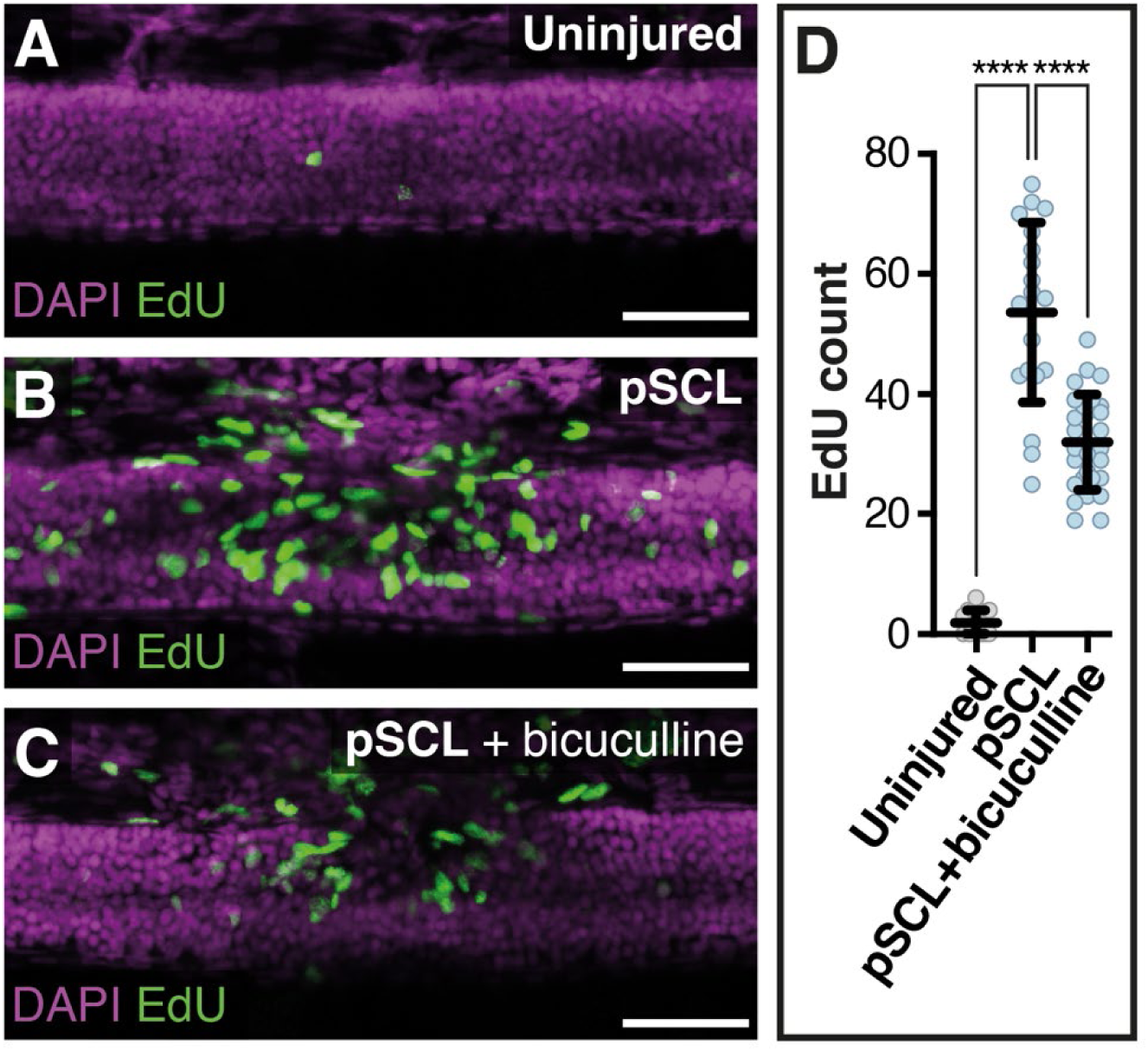
GABA_A_R signalling regulates progenitor activity in the regenerating spinal cord. (A–C) Representative EdU incorporation assays (green) at 5 dpf (48 h post-pSCL) in uninjured control (A), injured (B), and larvae treated with 25 μM bicuculline prior to injury (C). Scale bars, 40 μm. (D) Quantification of A–C.

**Figure S4:**
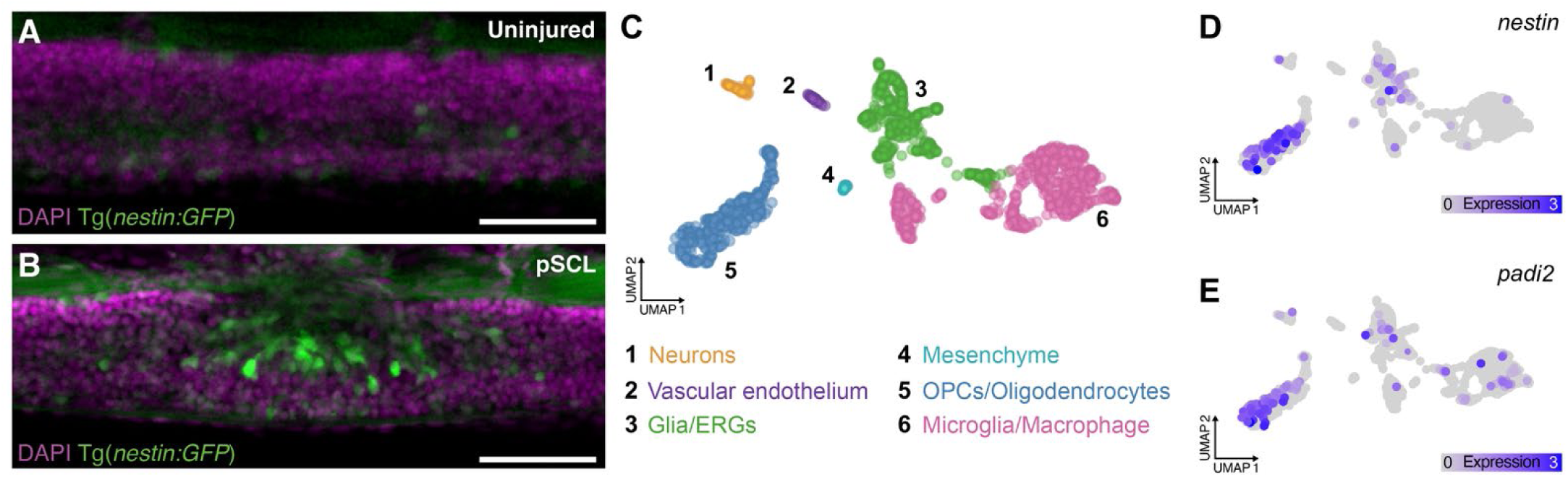
*nestin* expressing cells in the regenerating spinal cord also express *padi2*. (A–B) Sagittal views of *Tg(nestin:GFP)* larvae showing *nestin* reporter expression (green) and DAPI (magenta) in uninjured (A) and injured (B; 48 hpi) spinal cords. Scale bars, 50 μm. (C) UMAP projection of single-cell transcriptomic data from pooled larval trunks 1–2 days post-injury, filtered to exclude non-neural cell types. Colours indicate clusters representing major neural cell types, manually annotated from marker gene expression. (D–E) Feature plots showing *nestin* (D) and *padi2* (E) expression.

**Figure S5:**
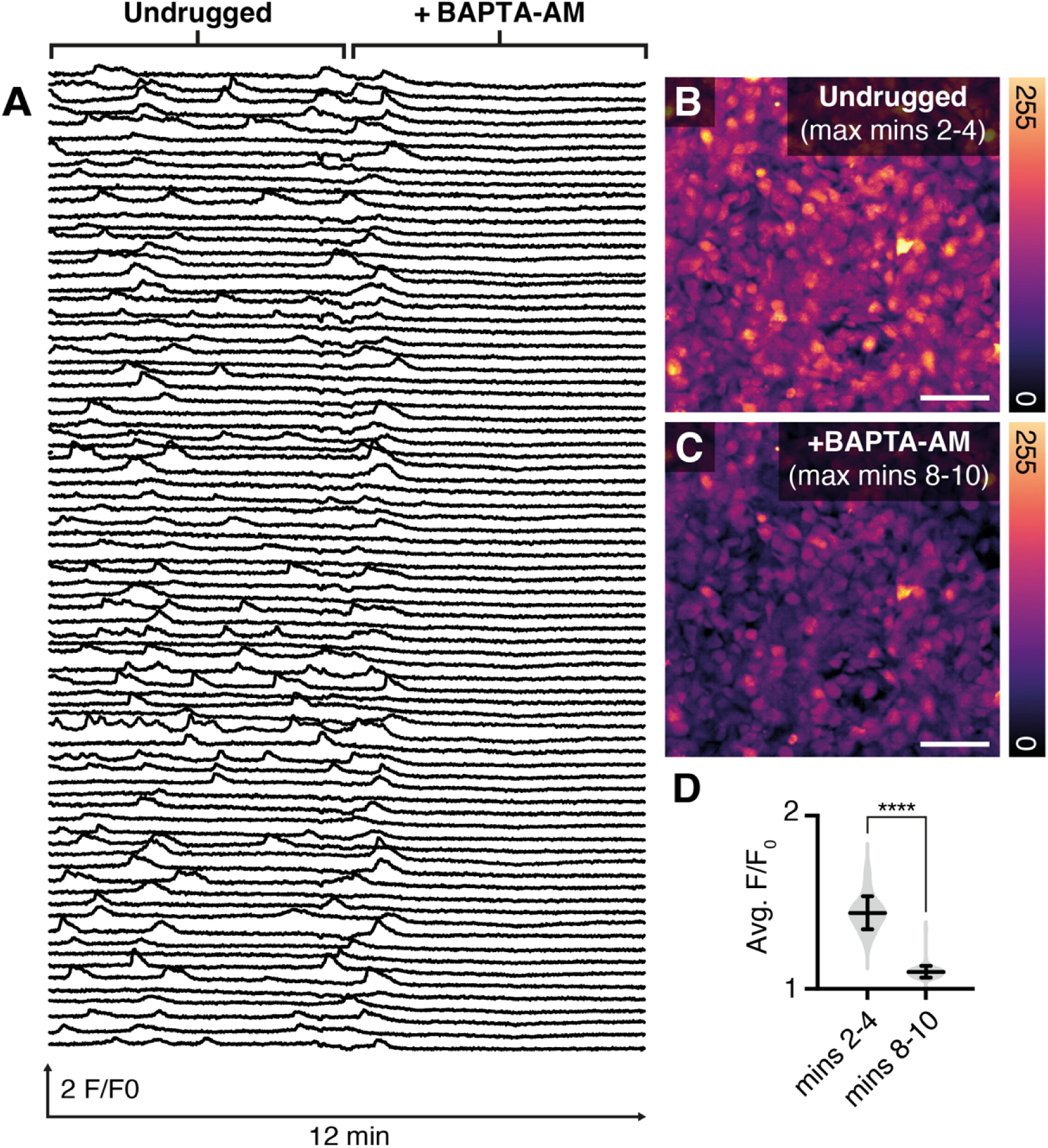
Spontaneous Ca2+ activity in confluent DAOY cells. (A) Normalised FLUO-4 fluorescence traces (F/F₀) in DAOY cells, showing normal activity (0–6 min) and reduced signal following BAPTA-AM treatment (6–12 min). (B–C) Maximum-intensity projections of FLUO-4 fluorescence over 2 min windows in untreated (B; 2–4 min) and BAPTA-AM–treated (C; 8–10 min) cells. (D) Quantification of average calcium signal (F/F₀) across time windows shown in (B–C).

**Figure S6:**
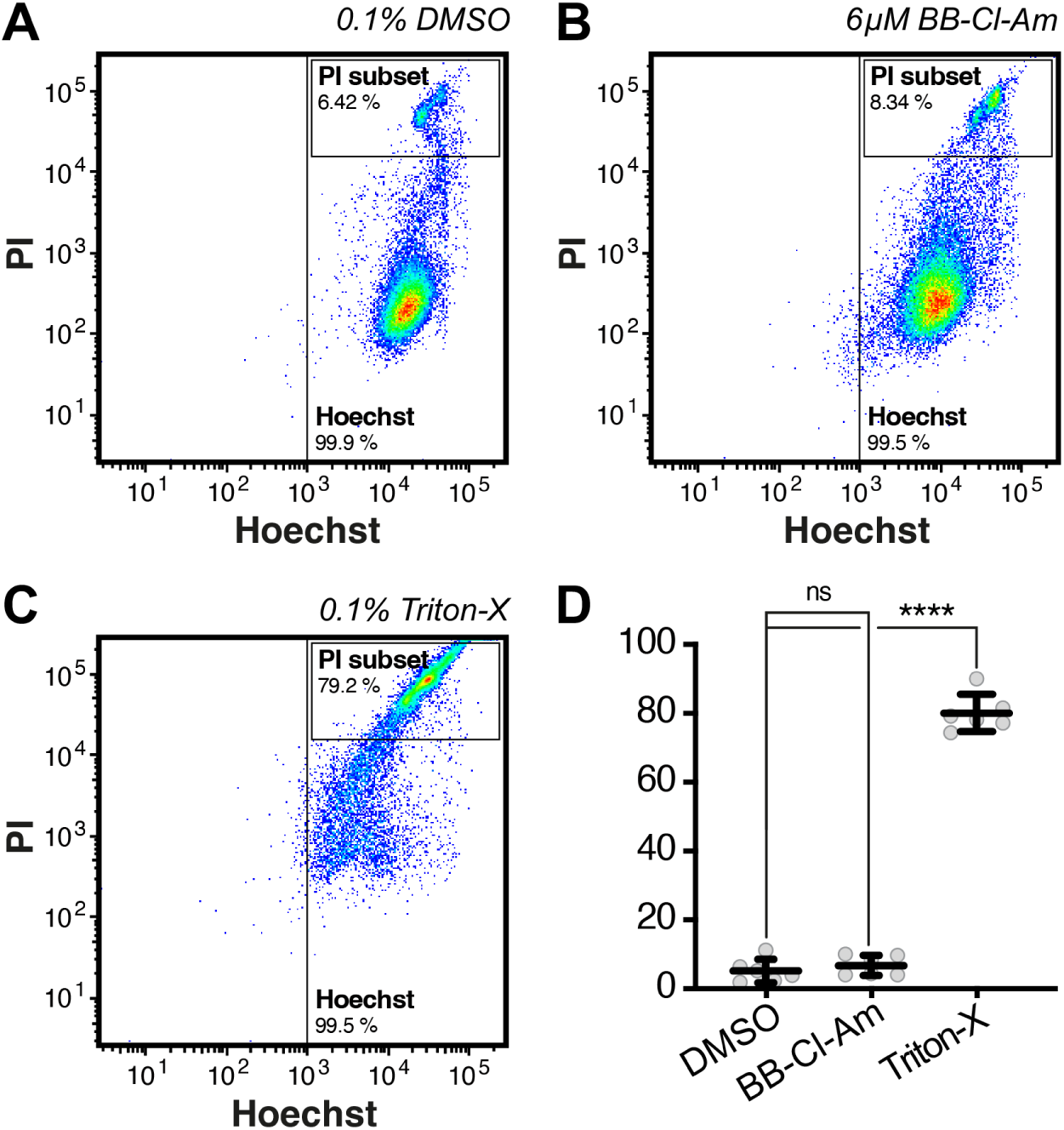
BB-Cl-amidine does not affect DAOY cell viability. (A–C) Flow cytometry plots showing propidium iodide (PI) versus Hoechst fluorescence in DAOY cells treated with 0.1% DMSO (A), 6 μM BB-Cl-amidine (B), or 0.1% Triton X (C, positive control). (D) Quantification of PI-labelled cells from conditions shown in (A–C).

**Figure S7:**
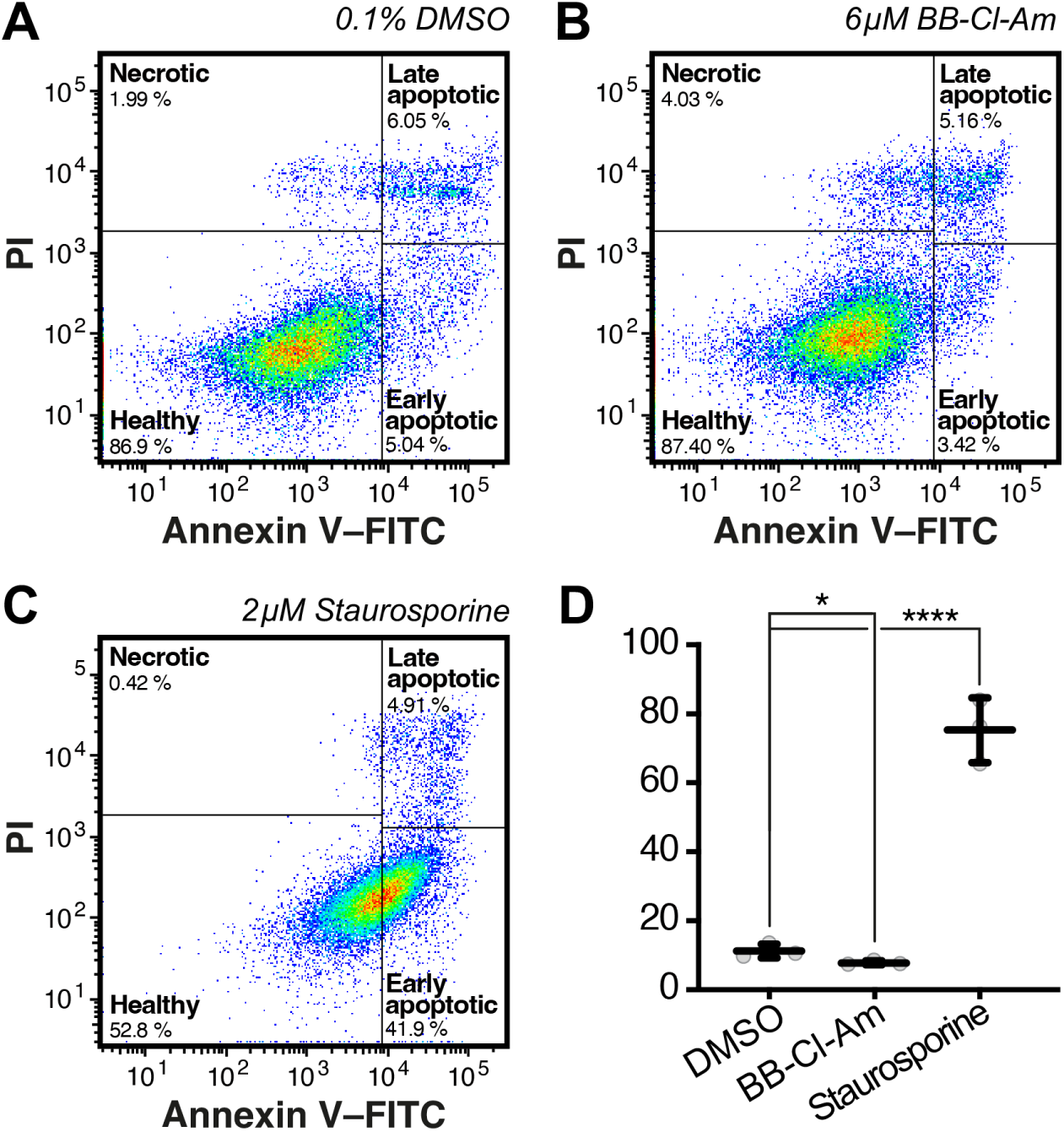
BB-Cl-amidine is not associated with increased apoptosis in DAOY cells. (A–C) Flow cytometry plots showing Annexin V–FITC versus propidium iodide (PI) staining in DAOY cells treated with 0.1% DMSO (A), 6 μM BB-Cl-amidine (B), or 2 μM staurosporine (C, positive control). (D) Quantification of early apoptotic cells (Annexin V⁺/PI⁻) from conditions shown in (A–C).

**Movie S1. Tracer particles injected into the CSF are retained in the central canal of both uninjured and injured larvae.** (A–B) Sagittal views of 5 dpf (2 dpi) larvae showing uninjured (A) and injured (pSCL, B) samples following injection of fluorescent tracer beads into the cerebrospinal fluid (CSF). Movies were acquired immediately after particle injection, showing retention of beads within the central canal in both experimental conditions.

**Movie S2. CSF circulation increases after injury.** (A–D) High-speed, high-resolution time-lapse recordings of circulating beads within the central canal of uninjured (A, B) and injured (C, D) 5 dpf (2 dpi) larvae. Top panels (A, C) show raw data, and bottom panels (B, D) show overlaid particle tracks colour-coded by mean track velocity.

**Movie S3. Motile ependymal cilia beat faster and elongate after injury.** (A–B) Time-lapse movie (A) and kymograph (B) of motile ependymal cilia within the central canal of an uninjured larva at 5 dpf. (C–D) Time-lapse movie (C) and kymograph (D) of motile ependymal cilia within the central canal of an injured larva at 5 dpf (2 dpi).

**Movie S4. The Tg(gad1b:Gal4) driver labels CSF-contacting neurons in the larval spinal cord.** Three-dimensional single time-point projection of a *Tg(gad1b:Gal4/UAS:GCaMP6s)*-expressing CSF-contacting neuron (CSF-cN) at 3 dpf (green). Approximately 1 nL of 70,000 MW Dextran Texas Red (magenta) was injected into the hindbrain ventricle immediately before imaging to visualise CSF.

**Movie S5. CSF-contacting neuron activity increases after injury.** (A–B) Representative high-speed time-lapse movies of *Tg(gad1b:Gal4/UAS:GCaMP6s)*-expressing CSF-cNs at 5 dpf (2 dpi), showing cells in the lesion-proximal region (A) and equivalent anatomical region of uninjured larvae (B).

**Movie S6. Basal ERG Ca²⁺ activity is low in the uninjured spinal cord.** (A, B) Time-matched single-channel timelapse movies of a representative uninjured *Tg(gfap:Gal4/UAS:GCaMP6s)* larva at 5 dpf. (A) Brightfield. (B) Maximum-intensity projection of GCaMP6s fluorescence spanning the full transverse spinal cord. (C) Quantification of (B).

**Movie S7. ERG Ca²⁺ activity increases after injury.** (A, B) Time-matched single-channel timelapse movies of a representative injured *Tg(gfap:Gal4/UAS:GCaMP6s)* larva at 5 dpf (2 dpi). (A) Brightfield. (B) Maximum-intensity projection of GCaMP6s fluorescence across the spinal cord showing elevated Ca²⁺ activity near the lesion. (C) Quantification of (B).

**Movie S8. CSF-contacting neuron activity regulates Ca²⁺ signalling in ERGs after injury.** (A, B) Time-matched single-channel timelapse movies of a representative injured *Tg(gfap:Gal4/UAS:GCaMP6s)* larva at 5 dpf (2 dpi) following F₀ CRISPR-mediated *pkd2l1* knockout. (A) Brightfield. (B) Maximum-intensity projection of GCaMP6s fluorescence showing reduced Ca²⁺ activity. (C) Quantification of (B).

**Movie S9. GABAₐR inhibition blocks ERG Ca²⁺ activity after injury.** (A, B) Time-matched single-channel timelapse movies of a representative injured *Tg(gfap:Gal4/UAS:GCaMP6s)* larva at 5 dpf (2 dpi) treated with 25 μM bicuculline. (A) Brightfield. (B) Maximum-intensity projection of GCaMP6s fluorescence showing suppressed Ca²⁺ activity. (C) Quantification of (B).

**Movie S10. Spontaneous Ca²⁺ activity in DAOY cells** (A, B) 12-minute time-lapse movie of confluent DAOY cultures loaded with the cell-permeable Ca²⁺ indicator Fluo-4 AM. BAPTA-AM was added at 6 min to establish a baseline for spontaneous Ca²⁺ activity.

## Bibliography

1. Navarro Negredo, P., Yeo, R.W., and Brunet, A. (2020). Aging and Rejuvenation of Neural Stem Cells and Their Niches. Cell Stem Cell 27, 202–223.

2. Urbán, N., Blomfield, I.M., and Guillemot, F. (2019). Quiescence of Adult Mammalian Neural Stem Cells: A Highly Regulated Rest. Neuron 104, 834–848.

3. Gonçalves, J.T., Schafer, S.T., and Gage, F.H. (2016). Adult Neurogenesis in the Hippocampus: From Stem Cells to Behavior. Cell 167, 897–914.

4. Vassal, M., Martins, F., Monteiro, B., Tambaro, S., Martinez-Murillo, R., and Rebelo, S. (2025). Emerging Pro-neurogenic Therapeutic Strategies for Neurodegenerative Diseases: A Review of Pre-clinical and Clinical Research. Mol Neurobiol 62, 46–76.

5. Wang, X., Zhou, R., Xiong, Y., Zhou, L., Yan, X., Wang, M., Li, F., Xie, C., Zhang, Y., Huang, Z., et al. (2021). Sequential fate-switches in stem-like cells drive the tumorigenic trajectory from human neural stem cells to malignant glioma. Cell Res 31, 684–702.

6. Azzarelli, R., Simons, B.D., and Philpott, A. (2018). The developmental origin of brain tumours: a cellular and molecular framework. Development 145, dev162693.

7. Wang, Y., Yang, J., Zheng, H., Tomasek, G.J., Zhang, P., McKeever, P.E., Lee, E.Y.H.P., and Zhu, Y. (2009). Expression of Mutant p53 Proteins Implicates a Lineage Relationship between Neural Stem Cells and Malignant Astrocytic Glioma in a Murine Model. Cancer Cell 15, 514–526.

8. Bardella, C., Al-Dalahmah, O., Krell, D., Brazauskas, P., Al-Qahtani, K., Tomkova, M., Adam, J., Serres, S., Lockstone, H., Freeman-Mills, L., et al. (2016). Expression of *Idh1*^R132H^ in the Murine Subventricular Zone Stem Cell Niche Recapitulates Features of Early Gliomagenesis. Cancer Cell 30, 578–594.

9. Kaslin, J., Ganz, J., Geffarth, M., Grandel, H., Hans, S., and Brand, M. (2009). Stem cells in the adult zebrafish cerebellum: initiation and maintenance of a novel stem cell niche. J Neurosci 29, 6142–6153.

10. Lindsey, B.W., Hall, Z.J., Heuzé, A., Joly, J.-S., Tropepe, V., and Kaslin, J. (2018). The role of neuro-epithelial-like and radial-glial stem and progenitor cells in development, plasticity, and repair. Progress in Neurobiology 170, 99–114.

11. Tendolkar, A., and Mokalled, M.H. (2025). Mechanisms underpinning spontaneous spinal cord regeneration. Development 152,

12. Tanaka, E.M., and Ferretti, P. (2009). Considering the evolution of regeneration in the central nervous system. Nat Rev Neurosci 10, 713–723.

13. Mokalled, M.H., Patra, C., Dickson, A.L., Endo, T., Stainier, D.Y., and Poss, K.D. (2016). Injury-induced ctgfa directs glial bridging and spinal cord regeneration in zebrafish. Science 354, 630–634.

14. Kang, J., Hu, J., Karra, R., Dickson, A.L., Tornini, V.A., Nachtrab, G., Gemberling, M., Goldman, J.A., Black, B.L., and Poss, K.D. (2016). Modulation of tissue repair by regeneration enhancer elements. Nature 532, 201–206.

15. Poss, K.D., and Tanaka, E.M. (2024). Hallmarks of regeneration. Cell Stem Cell 31, 1244–1261.

16. Lehtinen, Maria K., Zappaterra, Mauro W., Chen, X., Yang, Yawei J., Hill, A.D., Lun, M., Maynard, T., Gonzalez, D., Kim, S., Ye, P., et al. (2011). The Cerebrospinal Fluid Provides a Proliferative Niche for Neural Progenitor Cells. Neuron 69, 893–905.

17. Carrano, A., Zarco, N., Phillipps, J., Lara-Velazquez, M., Suarez-Meade, P., Norton, E.S., Chaichana, K.L., Quiñones-Hinojosa, A., Asmann, Y.W., and Guerrero-Cázares, H. (2021). Human Cerebrospinal Fluid Modulates Pathways Promoting Glioblastoma Malignancy. Frontiers in Oncology 11, 624145.

18. Ganz, J., Kaslin, J., Hochmann, S., Freudenreich, D., and Brand, M. (2010). Heterogeneity and Fgf dependence of adult neural progenitors in the zebrafish telencephalon. Glia 58, 1345–1363.

19. Petrik, D., Myoga, M.H., Grade, S., Gerkau, N.J., Pusch, M., Rose, C.R., Grothe, B., and Götz, M. (2018). Epithelial Sodium Channel Regulates Adult Neural Stem Cell Proliferation in a Flow-Dependent Manner. Cell Stem Cell 22, 865–878.

20. Djenoune, L., Mahamdeh, M., Truong, T.V., Nguyen, C.T., Fraser, S.E., Brueckner, M., Howard, J., and Yuan, S. (2023). Cilia function as calcium-mediated mechanosensors that instruct left-right asymmetry. Science 379, 71–78.

21. Thouvenin, O., Keiser, L., Cantaut-Belarif, Y., Carbo-Tano, M., Verweij, F., Jurisch-Yaksi, N., Bardet, P.L., van Niel, G., Gallaire, F., and Wyart, C. (2020). Origin and role of the cerebrospinal fluid bidirectional flow in the central canal. Elife 9, 47699.

22. Kramer-Zucker, A.G., Olale, F., Haycraft, C.J., Yoder, B.K., Schier, A.F., and Drummond, I.A. (2005). Cilia-driven fluid flow in the zebrafish pronephros, brain and Kupffer’s vesicle is required for normal organogenesis. Development 132, 1907–1921.

23. Grimes, D.T., Boswell, C.W., Morante, N.F., Henkelman, R.M., Burdine, R.D., and Ciruna, B. (2016). Zebrafish models of idiopathic scoliosis link cerebrospinal fluid flow defects to spine curvature. Science 352, 1341–1344.

24. Faubel, R., Westendorf, C., Bodenschatz, E., and Eichele, G. (2016). Cilia-based flow network in the brain ventricles. Science 353, 176–178.

25. Borovina, A., Superina, S., Voskas, D., and Ciruna, B. (2010). Vangl2 directs the posterior tilting and asymmetric localization of motile primary cilia. Nature Cell Biology 12, 407–412.

26. Huff, J. (2016). The Fast mode for ZEISS LSM 880 with Airyscan: high-speed confocal imaging with super-resolution and improved signal-to-noise ratio. Nature Methods 13, i–ii.

27. Vandestadt, C., Vanwalleghem, G.C., Khabooshan, M.A., Douek, A.M., Castillo, H.A., Li, M., Schulze, K., Don, E., Stamatis, S.A., Ratnadiwakara, M., et al. (2021). RNA-induced inflammation and migration of precursor neurons initiates neuronal circuit regeneration in zebrafish. Dev Cell 56, 2364–2380.e2368.

28. Kroll, F., Powell, G.T., Ghosh, M., Gestri, G., Antinucci, P., Hearn, T.J., Tunbak, H., Lim, S., Dennis, H.W., Fernandez, J.M., et al. (2021). A simple and effective F0 knockout method for rapid screening of behaviour and other complex phenotypes. eLife 10, e59683.

29. Hamimi, M., Khabooshan, M., Castillo, H.A., and Kaslin, J. (2019). Fluorescently Labeled TracrRNA Improves Work Flow and Facilitates Successful Genome Editing in Zebrafish. Zebrafish 16, 135–137.

30. Omori, Y., Zhao, C., Saras, A., Mukhopadhyay, S., Kim, W., Furukawa, T., Sengupta, P., Veraksa, A., and Malicki, J. (2008). elipsa is an early determinant of ciliogenesis that links the IFT particle to membrane-associated small GTPase Rab8. Nature Cell Biology 10, 437–444.

31. van Rooijen, E., Giles, R.H., Voest, E.E., van Rooijen, C., Schulte-Merker, S., and van Eeden, F.J. (2008). LRRC50, a Conserved Ciliary Protein Implicated in Polycystic Kidney Disease. Journal of the American Society of Nephrology 19, 1128–1138.

32. Shih, S.M., Engel, B.D., Kocabas, F., Bilyard, T., Gennerich, A., Marshall, W.F., and Yildiz, A. (2013). Intraflagellar transport drives flagellar surface motility. eLife 2, e00744.

33. Firestone, A.J., Weinger, J.S., Maldonado, M., Barlan, K., Langston, L.D., O’Donnell, M., Gelfand, V.I., Kapoor, T.M., and Chen, J.K. (2012). Small-molecule inhibitors of the AAA+ ATPase motor cytoplasmic dynein. Nature 484, 125–129.

34. Tsarouchas, T.M., Wehner, D., Cavone, L., Munir, T., Keatinge, M., Lambertus, M., Underhill, A., Barrett, T., Kassapis, E., Ogryzko, N., et al. (2018). Dynamic control of proinflammatory cytokines Il-1β and Tnf-α by macrophages in zebrafish spinal cord regeneration. Nature Communications 9, 4670.

35. Nelson, C.M., Lennon, V.A., Lee, H., Krug, R.G., Kamalova, A., Madigan, N.N., Clark, K.J., Windebank, A.J., and Henley, J.R. (2019). Glucocorticoids Target Ependymal Glia and Inhibit Repair of the Injured Spinal Cord. Frontiers in Cell and Developmental Biology 7, 00056.

36. Kyritsis, N., Kizil, C., Zocher, S., Kroehne, V., Kaslin, J., Freudenreich, D., Iltzsche, A., and Brand, M. (2012). Acute Inflammation Initiates the Regenerative Response in the Adult Zebrafish Brain. Science 338, 1353–1356.

37. Hui, S.P., Sheng, D.Z., Sugimoto, K., Gonzalez-Rajal, A., Nakagawa, S., Hesselson, D., and Kikuchi, K. (2017). Zebrafish Regulatory T Cells Mediate Organ-Specific Regenerative Programs. Developmental Cell 43, 659–672.e655.

38. de Sena-Tomás, C., Rebola Lameira, L., Rebocho da Costa, M., Naique Taborda, P., Laborde, A., Orger, M., de Oliveira, S., and Saúde, L. (2024). Neutrophil immune profile guides spinal cord regeneration in zebrafish. Brain, Behavior, and Immunity 120, 514–531.

39. Kamiya, Y., Fujisawa, T., Katsumata, M., Yasui, H., Suzuki, Y., Karayama, M., Hozumi, H., Furuhashi, K., Enomoto, N., Nakamura, Y., et al. (2020). Influenza A virus enhances ciliary activity and mucociliary clearance via TLR3 in airway epithelium. Respir Res 21, 282.

40. Tatematsu, M., Nishikawa, F., Seya, T., and Matsumoto, M. (2013). Toll-like receptor 3 recognizes incomplete stem structures in single-stranded viral RNA. Nature Communications 4, 1833.

41. Wyart, C., Carbo-Tano, M., Cantaut-Belarif, Y., Orts-Del’Immagine, A., and Böhm, U.L. (2023). Cerebrospinal fluid-contacting neurons: multimodal cells with diverse roles in the CNS. Nature Reviews Neuroscience 24, 540–556.

42. Sternberg, J.R., Prendergast, A.E., Brosse, L., Cantaut-Belarif, Y., Thouvenin, O., Orts-Del’Immagine, A., Castillo, L., Djenoune, L., Kurisu, S., McDearmid, J.R., et al. (2018). Pkd2l1 is required for mechanoception in cerebrospinal fluid-contacting neurons and maintenance of spine curvature. Nature Communications 9, 3804.

43. Prendergast, A.E., Jim, K.K., Marnas, H., Desban, L., Quan, F.B., Djenoune, L., Laghi, V., Hocquemiller, A., Lunsford, E.T., Roussel, J., et al. (2023). CSF-contacting neurons respond to Streptococcus pneumoniae and promote host survival during central nervous system infection. Curr Biol 33, 940–956.e910.

44. Orts-Del’Immagine, A., Seddik, R., Tell, F., Airault, C., Er-Raoui, G., Najimi, M., Trouslard, J., and Wanaverbecq, N. (2016). A single polycystic kidney disease 2-like 1 channel opening acts as a spike generator in cerebrospinal fluid-contacting neurons of adult mouse brainstem. Neuropharmacology 101, 549–565.

45. Chen, T.W., Wardill, T.J., Sun, Y., Pulver, S.R., Renninger, S.L., Baohan, A., Schreiter, E.R., Kerr, R.A., Orger, M.B., Jayaraman, V., et al. (2013). Ultrasensitive fluorescent proteins for imaging neuronal activity. Nature 499, 295–300.

46. Förster, D., Arnold-Ammer, I., Laurell, E., Barker, A.J., Fernandes, A.M., Finger-Baier, K., Filosa, A., Helmbrecht, T.O., Kölsch, Y., Kühn, E., et al. (2017). Genetic targeting and anatomical registration of neuronal populations in the zebrafish brain with a new set of BAC transgenic tools. Sci Rep 7, 5230.

47. England, S.J., Campbell, P.C., Banerjee, S., Swanson, A.J., and Lewis, K.E. (2017). Identification and Expression Analysis of the Complete Family of Zebrafish pkd Genes. Front Cell Dev Biol 5, 5.

48. Kanold, P.O., and Shatz, C.J. (2006). Subplate neurons regulate maturation of cortical inhibition and outcome of ocular dominance plasticity. Neuron 51, 627–638.

49. Ganguly, K., Schinder, A.F., Wong, S.T., and Poo, M. (2001). GABA itself promotes the developmental switch of neuronal GABAergic responses from excitation to inhibition. Cell 105, 521–532.

50. Gallo, V., Zhou, J.M., McBain, C.J., Wright, P., Knutson, P.L., and Armstrong, R.C. (1996). Oligodendrocyte progenitor cell proliferation and lineage progression are regulated by glutamate receptor-mediated K+ channel block. J Neurosci 16, 2659–2670.

51. LoTurco, J.J., Owens, D.F., Heath, M.J., Davis, M.B., and Kriegstein, A.R. (1995). GABA and glutamate depolarize cortical progenitor cells and inhibit DNA synthesis. Neuron 15, 1287–1298.

52. Nguyen, L., Rigo, J.-M., Rocher, V., Belachew, S., Malgrange, B., Rogister, B., Leprince, P., and Moonen, G. (2001). Neurotransmitters as early signals for central nervous system development. Cell and Tissue Research 305, 187–202.

53. Vitali, I., Fièvre, S., Telley, L., Oberst, P., Bariselli, S., Frangeul, L., Baumann, N., McMahon, J.J., Klingler, E., Bocchi, R., et al. (2018). Progenitor Hyperpolarization Regulates the Sequential Generation of Neuronal Subtypes in the Developing Neocortex. Cell 174, 1264–1276.e1215.

54. Haydar, T.F., Wang, F., Schwartz, M.L., and Rakic, P. (2000). Differential modulation of proliferation in the neocortical ventricular and subventricular zones. J Neurosci 20, 5764–5774.

55. Maric, D., Liu, Q.Y., Grant, G.M., Andreadis, J.D., Hu, Q., Chang, Y.H., Barker, J.L., Joseph, J., Stenger, D.A., and Ma, W. (2000). Functional ionotropic glutamate receptors emerge during terminal cell division and early neuronal differentiation of rat neuroepithelial cells. J Neurosci Res 61, 652–662.

56. Mayer, S., Chen, J., Velmeshev, D., Mayer, A., Eze, U.C., Bhaduri, A., Cunha, C.E., Jung, D., Arjun, A., Li, E., et al. (2019). Multimodal Single-Cell Analysis Reveals Physiological Maturation in the Developing Human Neocortex. Neuron 102, 143–158.e147.

57. Arjun McKinney, A., Petrova, R., and Panagiotakos, G. (2022). Calcium and activity-dependent signaling in the developing cerebral cortex. Development 149, dev198853.

58. Bernardos, R.L., and Raymond, P.A. (2006). GFAP transgenic zebrafish. Gene Expr Patterns 6, 1007–1013.

59. Stoeckel, M.E., Uhl-Bronner, S., Hugel, S., Veinante, P., Klein, M.J., Mutterer, J., Freund-Mercier, M.J., and Schlichter, R. (2003). Cerebrospinal fluid-contacting neurons in the rat spinal cord, a gamma-aminobutyric acidergic system expressing the P2X2 subunit of purinergic receptors, PSA-NCAM, and GAP-43 immunoreactivities: light and electron microscopic study. J Comp Neurol 457, 159–174.

60. Binor, E., and Heathcote, R.D. (2001). Development of GABA-immunoreactive neuron patterning in the spinal cord. Journal of Comparative Neurology 438, 1–11.

61. Vossenaar, E.R., Zendman, A.J.W., van Venrooij, W.J., and Pruijn, G.J.M. (2003). PAD, a growing family of citrullinating enzymes: genes, features and involvement in disease. BioEssays 25, 1106–1118.

62. Takahara, H., Okamoto, H., and Sugawara, K. (1986). Calcium-dependent Properties of Peptidylarginine Deiminase from Rabbit Skeletal Muscle. Agricultural and Biological Chemistry 50, 2899–2904.

63. Wang, Y., Li, M., Stadler, S., Correll, S., Li, P., Wang, D., Hayama, R., Leonelli, L., Han, H., Grigoryev, S.A., et al. (2009). Histone hypercitrullination mediates chromatin decondensation and neutrophil extracellular trap formation. J Cell Biol 184, 205–213.

64. Saiki, M., Watase, M., Matsubayashi, H., and Hidaka, Y. (2009). Recognition of the N-terminal histone H2A and H3 peptides by peptidylarginine deiminase IV. Protein Pept Lett 16, 1012–1016.

65. Cuthbert, G.L., Daujat, S., Snowden, A.W., Erdjument-Bromage, H., Hagiwara, T., Yamada, M., Schneider, R., Gregory, P.D., Tempst, P., Bannister, A.J., and Kouzarides, T. (2004). Histone deimination antagonizes arginine methylation. Cell 118, 545–553.

66. Zhang, X., Liu, X., Zhang, M., Li, T., Muth, A., Thompson, P.R., Coonrod, S.A., and Zhang, X. (2016). Peptidylarginine deiminase 1-catalyzed histone citrullination is essential for early embryo development. Sci Rep 6, 38727.

67. Singh, A.K., Khan, S., Moore, D., Andrews, S., and Christophorou, M.A. (2023). Transcriptomic analysis of PADI4 target genes during multi-lineage differentiation of embryonic stem cells. Philos Trans R Soc Lond B Biol Sci 378, 20220236.

68. Christophorou, M.A., Castelo-Branco, G., Halley-Stott, R.P., Oliveira, C.S., Loos, R., Radzisheuskaya, A., Mowen, K.A., Bertone, P., Silva, J.C., Zernicka-Goetz, M., et al. (2014). Citrullination regulates pluripotency and histone H1 binding to chromatin. Nature 507, 104–108.

69. Ballasy, N.N., Bering, E.A., Kokorudz, C., Radford, B.N., Zhao, X., Dean, W., and Hemberger, M. (2022). Padi2/3 Deficiency Alters the Epigenomic Landscape and Causes Premature Differentiation of Mouse Trophoblast Stem Cells. Cells 11, 2466.

70. Golenberg, N., Squirrell, J.M., Bennin, D.A., Rindy, J., Pistono, P.E., Eliceiri, K.W., Shelef, M.A., Kang, J., and Huttenlocher, A. (2020). Citrullination regulates wound responses and tissue regeneration in zebrafish. J Cell Biol 219, e201908164.

71. Bayin, N.S., Mizrak, D., Stephen, D.N., Lao, Z., Sims, P.A., and Joyner, A.L. (2021). Injury-induced ASCL1 expression orchestrates a transitory cell state required for repair of the neonatal cerebellum. Science Advances 7, eabj1598.

72. Kaslin, J., Kroehne, V., Ganz, J., Hans, S., and Brand, M. (2017). Distinct roles of neuroepithelial-like and radial glia-like progenitor cells in cerebellar regeneration. Development 144, 1462–1471.

73. Wojcinski, A., Lawton, A.K., Bayin, N.S., Lao, Z., Stephen, D.N., and Joyner, A.L. (2017). Cerebellar granule cell replenishment postinjury by adaptive reprogramming of Nestin+ progenitors. Nature Neuroscience 20, 1361–1370.

74. Yang, Z.J., Ellis, T., Markant, S.L., Read, T.A., Kessler, J.D., Bourboulas, M., Schüller, U., Machold, R., Fishell, G., Rowitch, D.H., et al. (2008). Medulloblastoma can be initiated by deletion of Patched in lineage-restricted progenitors or stem cells. Cancer Cell 14, 135–145.

75. Guo, D., Qu, Y., Yang, Y., and Yang, Z.J. (2020). Medulloblastoma cells resemble neuronal progenitors in their differentiation. Mol Cell Oncol 7, 1810514.

76. Jacobsen, P.F., Jenkyn, D.J., and Papadimitriou, J.M. (1985). Establishment of a human medulloblastoma cell line and its heterotransplantation into nude mice. J Neuropathol Exp Neurol 44, 472–485.

77. Shiau, C.E., Kaufman, Z., Meireles, A.M., and Talbot, W.S. (2015). Differential requirement for irf8 in formation of embryonic and adult macrophages in zebrafish. PLoS One 10, e0117513.

78. Bittman, K.S., and LoTurco, J.J. (1999). Differential Regulation of Connexin 26 and 43 in Murine Neocortical Precursors. Cerebral Cortex 9, 188–195.

79. LoTurco, J.J., Blanton, M.G., and Kriegstein, A.R. (1991). Initial expression and endogenous activation of NMDA channels in early neocortical development. J Neurosci 11, 792–799.

80. Marins, M., Xavier, A.L., Viana, N.B., Fortes, F.S., Fróes, M.M., and Menezes, J.R. (2009). Gap junctions are involved in cell migration in the early postnatal subventricular zone. Dev Neurobiol 69, 715–730.

81. Zhang, Y., Cao, L., Yan, H., Luo, Z., Chen, C., Shangguan, Z., Li, Q., Shi, X., Yang, L., Tan, W., et al. (2025). *pkd2l1* deletion inhibits the neurogenesis of cerebrospinal fluid-contacting neurons and impedes spinal cord injury repair. Cell Death Discovery 11, 194.

82. Böhm, U.L., Prendergast, A., Djenoune, L., Nunes Figueiredo, S., Gomez, J., Stokes, C., Kaiser, S., Suster, M., Kawakami, K., Charpentier, M., et al. (2016). CSF-contacting neurons regulate locomotion by relaying mechanical stimuli to spinal circuits. Nat Commun 7, 10866.

83. Nelson, Amanda M., Reddy, Sashank K., Ratliff, Tabetha S., Hossain, M.Z., Katseff, Adiya S., Zhu, Amadeus S., Chang, E., Resnik, Sydney R., Page, C., Kim, D., et al. (2015). dsRNA Released by Tissue Damage Activates TLR3 to Drive Skin Regeneration. Cell Stem Cell 17, 139–151.

84. Kim, D., Chen, R., Sheu, M., Kim, N., Kim, S., Islam, N., Wier, E.M., Wang, G., Li, A., Park, A., et al. (2019). Noncoding dsRNA induces retinoic acid synthesis to stimulate hair follicle regeneration via TLR3. Nature Communications 10, 2811.

85. Lin, Q., Fang, D., Fang, J., Ren, X., Yang, X., Wen, F., and Su, S.B. (2011). Impaired Wound Healing with Defective Expression of Chemokines and Recruitment of Myeloid Cells in TLR3-Deficient Mice. The Journal of Immunology 186, 3710–3717.

86. Saraswathy, V.M., Zhou, L., and Mokalled, M.H. (2024). Single-cell analysis of innate spinal cord regeneration identifies intersecting modes of neuronal repair. Nature Communications 15, 6808.

87. Venkataramani, V., Tanev, D.I., Strahle, C., Studier-Fischer, A., Fankhauser, L., Kessler, T., Körber, C., Kardorff, M., Ratliff, M., Xie, R., et al. (2019). Glutamatergic synaptic input to glioma cells drives brain tumour progression. Nature 573, 532–538.

88. Venkatesh, Humsa S., Johung, Tessa B., Caretti, V., Noll, A., Tang, Y., Nagaraja, S., Gibson, Erin M., Mount, Christopher W., Polepalli, J., Mitra, Siddhartha S., et al. (2015). Neuronal Activity Promotes Glioma Growth through Neuroligin-3 Secretion. Cell 161, 803–816.

89. Chigurupati, S., Venkataraman, R., Barrera, D., Naganathan, A., Madan, M., Paul, L., Pattisapu, J.V., Kyriazis, G.A., Sugaya, K., Bushnev, S., et al. (2010). Receptor Channel TRPC6 Is a Key Mediator of Notch-Driven Glioblastoma Growth and Invasiveness. Cancer Research 70, 418–427.

90. Visa, A., Sallán, M.C., Maiques, O., Alza, L., Talavera, E., López-Ortega, R., Santacana, M., Herreros, J., and Cantí, C. (2019). T-Type Cav3.1 Channels Mediate Progression and Chemotherapeutic Resistance in Glioblastoma. Cancer Research 79, 1857–1868.

91. Liu, G., Zhang, H., Chen, S., Gao, J., Zhao, H., Dong, Y., Liu, C., Wei, X., Li, T., Lu, C., et al. (2025). Mitochondrial Calcium Uniporter Links Acetyl-CoA Metabolism and H3K27 Acetylation to Maintain Glioblastoma Stem Cells. Cancer Research 85, 3416–3434.

92. Cárdenas, C., Müller, M., McNeal, A., Lovy, A., Jaňa, F., Bustos, G., Urra, F., Smith, N., Molgó, J., Diehl, J.A., et al. (2016). Selective Vulnerability of Cancer Cells by Inhibition of Ca^2+^ Transfer from Endoplasmic Reticulum to Mitochondria. Cell Reports 14, 2313–2324.

93. Barron, T., Yalçın, B., Su, M., Byun, Y.G., Gavish, A., Shamardani, K., Xu, H., Ni, L., Soni, N., Mehta, V., et al. (2025). GABAergic neuron-to-glioma synapses in diffuse midline gliomas. Nature 639, 1060–1068.

94. Tillotson, J., Aryal, B., Lai, L., Beaver, J.A., and Rao, V.A. (2023). Differential Protein Citrullination in Human ER– and ER+ Tumor and Adjacent Healthy Breast Tissue. Biochemistry 62, 893–898.

95. Xue, T., Liu, X., Song, C., Fei, S., Gu, J., Han, Y., Xing, J., Liu, X., Liang, F., Thompson, P.R., and Zhang, X. (2025). Citrullination of AKT2 Catalyzed by PAD1 Facilitates the Maintenance of Stemness Characteristics of Ovarian Cancer Stem-Like Cells in Ovarian Cancer. Adv Sci (Weinh) e01014.

96. Wang, L., Song, G., Zhang, X., Feng, T., Pan, J., Chen, W., Yang, M., Bai, X., Pang, Y., Yu, J., et al. (2017). PADI2-Mediated Citrullination Promotes Prostate Cancer Progression. Cancer Res 77, 5755–5768.

97. Albano, C., Biolatti, M., Mazibrada, J., Pasquero, S., Gugliesi, F., Lo Cigno, I., Calati, F., Bajetto, G., Riva, G., Griffante, G., et al. (2024). PAD-mediated citrullination is a novel candidate diagnostic marker and druggable target for HPV-associated cervical cancer. Front Cell Infect Microbiol 14, 1359367.

98. Lin, H.Y., Yu, C.C., Chi, C.L., Wei, C.K., Yin, W.Y., Tseng, C.E., and Li, S.C. (2023). Peptidylarginine Deiminase Type 2 Predicts Tumor Progression and Poor Prognosis in Patients with Curatively Resected Biliary Tract Cancer. Cancers (Basel) 15, 4131.

99. Chang, X., Chai, Z., Zou, J., Wang, H., Wang, Y., Zheng, Y., Wu, H., and Liu, C. (2019). PADI3 induces cell cycle arrest via the Sirt2/AKT/p21 pathway and acts as a tumor suppressor gene in colon cancer. Cancer Biol Med 16, 729–742.

100. Westerfield, M. (2007). The zebrafish book: A guide for the laboratory use of zebrafish Danio (“Brachydanio Rerio”) (University of Oregon).

101. Mu, Y., Bennett, D.V., Rubinov, M., Narayan, S., Yang, C.-T., Tanimoto, M., Mensh, B.D., Looger, L.L., and Ahrens, M.B. (2019). Glia Accumulate Evidence that Actions Are Futile and Suppress Unsuccessful Behavior. Cell 178, 27–43.e19.

102. Miles, L.B., Calcinotto, V., Oveissi, S., Serrano, R.J., Sonntag, C., Mulia, O., Lee, C., and Bryson-Richardson, R.J. (2024). CRIMP: a CRISPR/Cas9 insertional mutagenesis protocol and toolkit. Nature Communications 15, 5011.

103. Crossman, S.H., Khabooshan, M.A., Stamatis, S.A., Vandestadt, C., and Kaslin, J. (2024). Mechanical Ablation of Larval Zebrafish Spinal Cord. Methods Mol Biol 2746, 47–56.

104. Peterson, S.M., and Freeman, J.L. (2009). RNA isolation from embryonic zebrafish and cDNA synthesis for gene expression analysis. J Vis Exp

105. Susaki, E.A., Tainaka, K., Perrin, D., Yukinaga, H., Kuno, A., and Ueda, H.R. (2015). Advanced CUBIC protocols for whole-brain and whole-body clearing and imaging. Nature Protocols 10, 1709–1727.

106. Póvoa, V., Rebelo de Almeida, C., Maia-Gil, M., Sobral, D., Domingues, M., Martinez-Lopez, M., de Almeida Fuzeta, M., Silva, C., Grosso, A.R., and Fior, R. (2021). Innate immune evasion revealed in a colorectal zebrafish xenograft model. Nature Communications 12, 1156.

107. Pylvänäinen, J.W., Laine, R.F., Saraiva, B.M.S., Ghimire, S., Follain, G., Henriques, R., and Jacquemet, G. (2023). Fast4DReg - fast registration of 4D microscopy datasets. J Cell Sci 136, jcs260728.

108. Hao, Y., Stuart, T., Kowalski, M.H., Choudhary, S., Hoffman, P., Hartman, A., Srivastava, A., Molla, G., Madad, S., Fernandez-Granda, C., and Satija, R. (2024). Dictionary learning for integrative, multimodal and scalable single-cell analysis. Nature Biotechnology 42, 293–304.

